# Green-gradient based canopy segmentation: A multipurpose image mining model with potential use in crop phenotyping and canopy studies

**DOI:** 10.1101/241786

**Authors:** Abbas Haghshenas, Yahya Emam

## Abstract

Efficient quantification of the sophisticated shading patterns inside the 3D vegetation canopies may improve our understanding of canopy functions and status, which is possible now more than ever, thanks to the high-throughput phenotyping (HTP) platforms. In order to evaluate the option of quantitative characterization of shading patterns, a simple image mining technique named “*green-gradient based canopy segmentation model (GSM)*” was developed based on the relative variations in the level of RGB triplets under different illuminations. For this purpose, an archive of ground-based nadir images of heterogeneous wheat canopies (cultivar mixtures) was analyzed. The images were taken from experimental plots of a two-year field experiment conducted during 2014-15 and 2015-16 growing seasons in the semi-arid region of southern Iran. In GSM, the vegetation pixels were categorized into the maximum possible number of 255 groups based on their green levels. Subsequently, mean red and mean blue levels of each group were calculated and plotted against the green levels. It is evidenced that the yielded graph could be readily used for (i) identifying and characterizing canopies even as simple as one or two equation(s); (ii) classification of canopy pixels in accordance with the degree of exposure to sunlight; and (iii) accurately prediction of various quantitative properties of canopy including canopy coverage (CC), Normalized difference vegetation index (NDVI), canopy temperature, and also precise classification of experimental plots based on the qualitative characteristics such as subjecting to water and cold stresses, date of imaging, and time of irrigation. It seems that the introduced model may provide a multipurpose HTP platform and open new windows to canopy studies.

## 1. Introduction

In plant communities, a theoretical flat (2D) arrangement of a single unshaded layer of green leaves would potentially provide the light conditions for photosynthesis; though, such conceptually integrated light-exposed surface is actually distributed in a sophisticated manner over a three dimensional structure, which inevitably casts a shadow on itself. Moreover, the shading pattern of canopies shows large tempo-spatial variations e.g. due to time, season, plant growth stage, altering growth behaviors, biotic and abiotic stresses, etc.; which makes it even more challenging to approach coincidentally a comprehensive, efficient, and easily applicable characterizing framework.

Attempts to respond the intensified needs for a modern way of observation in plant science and agriculture, has led to the era of high-throughput phenotyping (HTP), i.e. the ubiquitous utilization of the remotely quantifying platforms for a deeper sensation of vegetation canopies. With respect to the key roles predicted for HTP, e.g. in mapping the genotype to phenotype (Chen *et al.*, 2014), its integration with genomics is even described as the potentially second green revolution (Tanger *et al.*, 2017; Crain *et al.*, 2018). Such a promising perspective is basically originated from the significant capacities of HTP for data collecting and processing (e.g. see Cabrera-Bosquet *et al.*, 2012; Araus and Cairns, 2014; Zhang *et al.*, 2017). Besides the advantages of HTP in producing huge amounts of data, the platforms efficiency is rarely considered (see Araus *et al.*, 2018); which may be defined using the concept of the balance between the complexity of the sensing and/or computation units on one hand, and the resulted achievements on the other hand. For instance, the method efficiency is expected to be higher where adequate and reliable information about crop canopy is easily achieved using image processing of a common digital image, compared with the case of using multiple types of sensors and/or sophisticated computation models, with a more or less similar outputs.

The high spectral overlap between the electromagnetic radiation of visible light and photosynthetically active radiation (PAR, i.e. about 400-700 nm wavelengths; McCree, 1972), provides the opportunity of using digital cameras in crop phenotyping as both reliable and readily available sensors of remote sensing. Interestingly, most of the current low-cost approaches for crop phenotyping are based on exploitation of the possibilities opened by RGB images (Araus *et al.*, 2018). Employing commercial digital cameras along with most often basic image processing techniques, an increasing number of studies have reported robust correlations between various image-derived indices and crop criteria including nitrogen status (Li *et al.*, 2010; Wang *et al.*, 2013; Lee and Lee, 2013), crop growth (Sakamoto *et al.*, 2012; Lee and Lee, 2013), chlorophyll content (Hunt Jr *et al.*, 2013), leaf area (Easlon and Bloom, 2014), leaf angle (Zou *et al.*, 2014), selection criteria for breeding programs (Casadesús *et al.*, 2007), etc.

Following the improvements in the 3D visualization technology, a reasonable strategy has been utilized for crop HTP is using models based on 3D reconstruction of single plants or canopies (e.g. see Burgess *et al.*, 2017; Gibbs *et al.*, 2018; Artzet *et al.*, 2019; Li *et al.*, 2019; Paulus *et al.*, 2019; and Shi *et al.*, 2019); which may be the ultimate level of image-based phenotyping. Indeed, in such developed approaches, a comprehensive representation of the plant/ or canopy volume is provided by digitalizing almost every point of plant as fine voxels. Therefore, it is expected that the required data for analyzing almost every 3D feature of the canopy e.g. spatial gap percentage, stem or leaf dimensions and angle, and even more complex indices will be available for researchers. However, utilizing the 3D-based models faces various technical challenges; as for instance, multi-directional imaging and usually sophisticated computation algorithms are needed. On the other hand, at least where the purpose is not determining the exact 3D features themselves –e.g. leaf shape or spatial arrangement-, it seems that the single 2D nadir images may also indirectly provide valuable data for the canopy 3D structure. The key may be analyzing the shading pattern within the canopy, which is the direct result of the interaction between light and the 3D arrangement of canopy.

As in many other fields of empirical sciences, sampling for statistical analyses is a fundamental factor in crop/plant physiological assessments; where probably the most significant power of digital cameras lies. For instance, a typical 8-megapixel color image taken from a low height above a crop canopy with 80% coverage, may provide a robust physiological sample of 6 million pixels. In addition to spatial resolution, high color sensibility, e.g. the ability of discriminating over 16 million – or 256^3^-colors per pixel in a common 24-bit color depth, has made it possible to record the considerable number of 256 different mono-color intensities of light reflected in either red, green, and blue bands per each single pixel (it is defined as a value between 0 to 255, i.e. zero to maximum intensity of light sensed in a single band). Processing the certainly adequate dataset provided by such a beneficial light sampling tool, requires subsequently more efficient data management, computational approaches, and powerful interpretation of results (Araus and Cairns, 2014) in order to improve our understanding of crop phenotype. Accordingly, it seems that a minor part of the computation capacity provided by RGB images has been explored.

Image mining of an archive of ground-based nadir images taken from the experimental plots of a field study consisted of monocultures and mixtures of early-to middle-ripening wheat cultivars, an unexpected observation was made using the statistics of the RGB triplets, based on which the present study was designed. As an option for evaluating the shading pattern within canopy, we were seeking a method to categorize the vegetation parts of the image based on their exposure to the sunlight. Therefore, the pattern of relative variations in the levels of the R, G, and B colors was evaluated from the darkest vegetation pixels towards the lightest ones in the image. To implement this idea, the vegetation pixels were categorized into the maximum possible number of 255 groups based on their green levels (i.e. in the range of 1-255; RGB color system). Subsequently, mean red and mean blue levels of each group were calculated, and the total sets of the results were plotted against the green ascending sequence. It was observed that sets of the red and blue mean values yielded two smooth upward exponential shaped curves, among which the blue one had apparently a higher degree of curvature compared with red. Since the set of ascending green values was used for classification of vegetation pixels, the approach was named *Green-gradient based canopy segmentation model* (GSM). It is notable that, although other color spaces such as HIS (e.g. see Cue *et al.*, 2010), HSV (Hamuda *et al.*, 2017), or LAB (Reza *et al.*, 2019), are available for image processing, the present study was conducted in the RGB color space, because the main idea of GSM was developed based on comparing variations in the level of red, green and blue colors, and RGB is the only color space that directly represents the color as the three distinct values of R, G, and B triplets, so makes such comparisons possible.

Considering the simplicity of the model in both implementation and interpretation, it appeared that GSM might be developed as a relatively efficient computation platform for RGB-based crop phenotyping. Therefore, the purpose of the present study was assessing the option of using GSM as (i) a tool for evaluating the shading pattern within canopy using single 2D ground-based nadir images, and also (ii) as a source of efficacious vegetation criteria in high-throughput phenotyping approaches.

## 2. Materials and Methods

### 2.1. Field experiment

The two-year field experiment was conducted at the research field of School of Agriculture, Shiraz University, Iran (29°73’ N latitude and 52°59’ E longitude at an altitude of 1,810 masl) during the 2014-15 and 2015-16 growing seasons. The experiment consisted of 90 (2×2 meter) plots including every 15 monocultures and mixtures of four early-to middle-ripening wheat cultivars grown under two normal and post-anthesis deficit irrigation conditions with 3 replicates. The experimental design was Randomized Complete Block Design (RCBD) in a lattice arrangement with 1 meter distances between the plots. The early-to middle-ripening wheat cultivars used were Chamran (1), Sirvan (2), Pishtaz (3), and Shiraz (4), respectively. Before sowing on flat beds, seeds of the cultivars were mixed with equal ratios based on their 1000-grain weights, germination percentages, and the fixed planting density of 450 plants per square meter. The planting dates in the first and second growing seasons were November 20 and November 5, respectively. Depending on soil test, only nitrogen fertilizer (urea) was applied in 3 equal splits i.e. at sowing, early tillering, and anthesis, with a total amount of 150 kg N/ha. No chemical was used, and weeding was done by hand.

Irrigation interval was 10 days based on local practices, and the amount of irrigation water was estimated using the Fao-56 Penman-Monteith model with local corrected coefficients which was reduced to 50% of evapo-transpirational demand from the first irrigation after anthesis (see Haghshenas and Emam, 2019D). Late in the season, plants were harvested from the center of plots (equal to the overall row length of 3 m, or 0.6 m^2^ per plot) and grain yield was calculated using a laboratory thresher and weighing.

### 2.2. Imaging

Nadir images of the experimental plots were taken regularly during both growing seasons from 150 cm above the soil surface. We used a common commercial digital camera (Canon PowerShot SX100 *IS*, Japan, Tokyo) which was set to auto mode and the maximum imaging resolution of 8.0 megapixels. Among the image archive, datasets of 10 imaging dates were selected for GSM analyses, depended on crop growth stage (mostly during post-anthesis period), occurrence of environmental stresses, or coincidence with other in-field measurements (Datasets A to J, Table 1). Due to occurrence of a moderate cold stress in the second year, the data of respective imaging date (Dataset D) was also used for evaluating the potential effects of stress on the GSM output.

**Table 1.** Properties of the image datasets used in GSM analyses.

| Data set | Imaging date | Azimuth<br>(initial-end) | (initial-Zenith<br>(initial-end)) | Azimuth<br>at solar<br>noon | Zenith at<br>solar noon | DAS | Growth stage | Spike | No.<br>images<br>(plots) | Analyses |  |  |  |  |  |  |  |
| --- | --- | --- | --- | --- | --- | --- | --- | --- | --- | --- | --- | --- | --- | --- | --- | --- | --- |
|  |  |  |  |  |  |  |  |  |  | CC | Canopy<br>temperature | NDVI | Grain yield | Water<br>stress | Cold stress | Irrigation<br>event | Imaging<br>date |
| A | 4/25/2015 | 261.70 - 266.7 | 40.55 - 33.00 | 180.00 | 73.40 | 156 | Late booting to early spike emergence | Not emerged | 90 | ✓ | ✓ | - | ✓ | - | ✓ | - | ✓ |
| B | 5/29/2015 | 256.67 - 260.84 | 65.48 - 61.23 | 180.00 | 81.86 | 190 | Milky to soft dough | Emerged | 56 | ✓ | ✓ | - | ✓ | ✓ | ✓ | ✓ | ✓ |
| C | 6/1/2015 | 242.58 - 255.1 | 74.96 - 67.86 | 180.00 | 82.30 | 193 | Milky to soft dough | Emerged | 90 | ✓ | ✓ | - | ✓ | ✓ | ✓ | ✓ | ✓ |
| D | 4/27/2016 | N/A | N/A | N/A | N/A | 174 | Late booting to early spike emergence | Not emerged | 90 | - | - | - | ✓ | - | ✓ | - | ✓ |
| E | 5/11/2016 | 257.68 - 263.64 | 56.32 - 48.83 | 180.00 | 78.30 | 188 | Late anthesis | Emerged | 90 | ✓ | ✓ | ✓ | ✓ | - | ✓ | ✓ | ✓ |
| F | 5/14/2016 | 268.13 - 272.45 | 43.72 - 36.13 | 180.01 | 79.04 | 191 | Milky stage | Emerged | 90 | ✓ | ✓ | ✓ | ✓ | - | ✓ | ✓ | ✓ |
| G | 5/20/2016 | 239.95 - 252.39 | 72.66 - 65.7 | 180.00 | 80.37 | 197 | Milky to soft dough | Emerged | 90 | ✓ | ✓ | ✓ | ✓ | ✓ | ✓ | ✓ | ✓ |
| H | 5/23/2016 | 260.21 - 265.92 | 59.68 - 52.14 | 180.01 | 80.96 | 200 | Milky to soft dough | Emerged | 90 | ✓ | ✓ | ✓ | ✓ | ✓ | ✓ | ✓ | ✓ |
| I | 5/31/2016 | 246.96 - 257.48 | 72.97 - 65.73 | 180.00 | 82.26 | 208 | Soft dough | Emerged | 90 | ✓ | ✓ | ✓ | ✓ | ✓ | ✓ | ✓ | ✓ |
| J | 6/3/2016 | 263.92 - 268.97 | 59.56 - 51.98 | 180.01 | 82.65 | 211 | Soft dough | Emerged | 90 | ✓ | ✓ | ✓ | ✓ | ✓ | ✓ | ✓ | ✓ |

Images of datasets A, B, and C were taken during the first season and the other datasets were selected from the archive of the second season. DAS: days after sowing; CC: canopy cover; NDVI: Normalized difference vegetation index. In the dataset B, images of 34 plots were missed. Range of solar azimuth and zenith at imaging time were not used in the calculations of the present study. "✓" shows that the respective dataset was used in the intended analysis. It is expected that appearance of spikes in the image could significantly affect the analysis.

### 2.3. Image processing and data analyses

#### 2.3.1. Segmentations types 1 and 2

Image processing was carried out using an exclusive MATLAB code (a user-friendly version of the code is published as a reproducible compute capsule on Code Ocean at https://doi.org/10.24433/CO.4355649.v1). Following a common segmentation of each image into vegetation and background parts i.e. segmentation type 1 or ST_1_ using the thresholding formula of G-R>0 (Wang *et al.*, 2016; G and R stand for green and red color values in RGB color system, respectively), pixels of vegetation parts were categorized into the maximum number of 255 groups, based on their similar green values (segmentation type 2, ST_2_; RGB color system). Then, the averaged red and averaged blue values of each group were calculated, and subsequently their separate trends were plotted against the green sequence.

In ST_2_, level of green color was selected as the basic exposure criterion, since although we had red, blue, grayscale, and other criteria as alternatives, the green band has obviously the most rigorous reflection from vegetation, which is expected to provide the most robust signal and also determines the exposure status more accurately, compared with the red and blue colors. In addition, it is directly a record of the light intensity in a given band captured by the sensor, despite the other formula-based indices e.g. the synthesized grayscale. Below is the mathematical representation of GSM i.e. ST_2_:

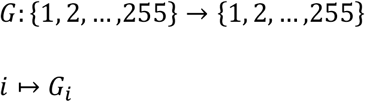

*where G_i_ is the green level of vegetation pixels (in RGB color system)*

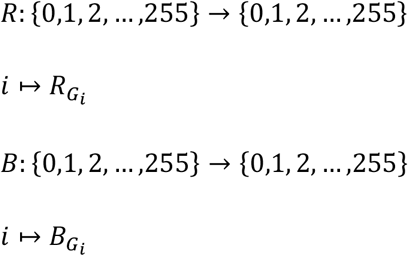

*where* 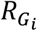 *and* 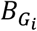 *are the red and blue levels of pixels with G_i_* = *i*.

*Let*:

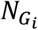 := *Number of pixels with G_i_* = *i*.
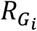 := *Red level of pixels with G_i_* = *i*.
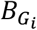 := *Blue level of pixels with G_i_* = *i*.

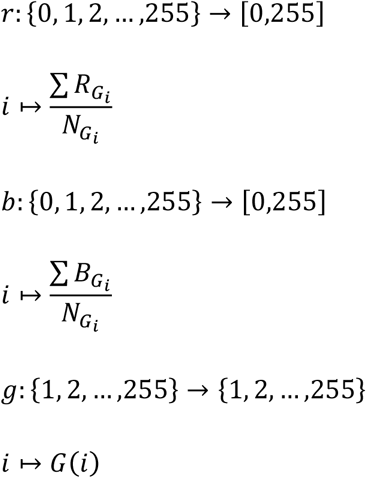

#### 2.3.2. Form of GSM red and blue curves

For evaluating shape of GSM curves, simple exponential equations were fitted to the red and blue curves as shown below:

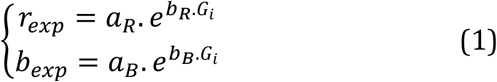

where *r*_*exp*_ and *b*_*exp*_ are the exponential equations of the GSM red and blue curves, and *a*_*R*_, *a*_*B*_, *b*_*R*_, *b*_*B*_ are the exponential coefficients of GSM red and blue curves.

#### 2.3.3. Segmentation type 3

Comparing the local slopes (differentiations) of the GSM red and blue curves with the slope of the green line i.e. equals 1, five apparent parts (classes) were distinguished on each of the red and blue curves, based on which the curves were segmented (Fig. 1). This curve segmentation was carried out using an exclusive algorithm (Fig. 2) and named “segmentation type 3 (ST_3_)” for avoiding confusion with the previous types of segmentation. Subsequently, the vegetation parts of the respective images were segmented based on ST_3_ thresholds and evaluated. For this purpose, the extreme green levels (thresholds) of each ST_3_ curve segment were used as inputs for image segmentation. In order to monitoring the calculation processes and results carefully, differentiations of GSM red and blue curves were checked manually with the results of Origin Pro 8 software (OriginLab, Northampton, MA). It is notable that ST_3_ was only carried out for images of Dataset A as examples. At this growth stage, wheat stand has a sufficiently dense canopy, most leaves are fully expanded and still thoroughly green, and also spikes are not emerged yet. Therefore, the segmentation method evaluated for such a typical canopy may be also generalized to a broader spectrum of vegetation stands.

**Figure 1.**
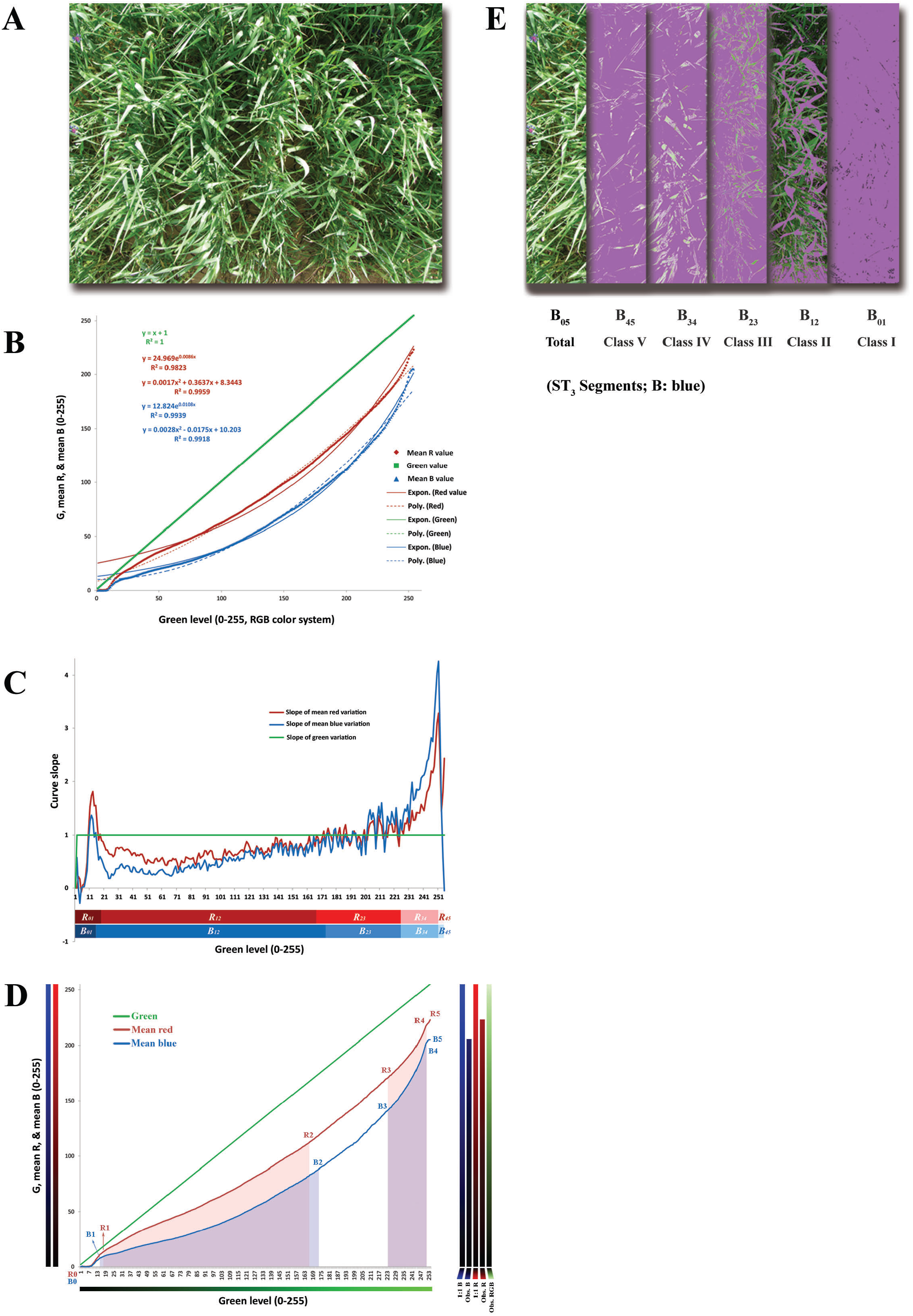
Illustration of green-gradient based canopy segmentation (GSM) and the GSM-curve derived segmentation type 3 (ST_3_). **(A)** the original image taken from monoculture ofthel st wheat cultivar. All ofthe graphs in other parts are resulted from processing this image. **(B)** Canopy GSM graph. The trend lines ofthe best polynomial and exponential equations fitted on the red and blue curves are shown. **(C)** The relative variations in local slopes ofthe red and blue GSM curves, compared to the green line. ST_3_ is developed based on this graph; see the horizontal red and blue bars below the graph, which are divided into 5 ST_3_ classes. **(D)** Segmentation type 3 (i.e. the GSM-curve segmentation based on status ofvariations in local slopes compared to the green trend). The set of five vertical color bars at the left side show the comparative theoretical and actual color variations. The color bars are constructed using the real outputs of GSM, as the “RGB” bar is made using the overall mean of red, green, and blue values at each green level; the “Obs. R” and “Obs. B” bars indicate the actual attenuating trend ofred and blue colors in the GSM graph, respectively; and the “1:1 R” and “1:1 B” bars show the conceptual attenuating trends ofthe red and blue colors, ifthey had declined linearly as same as the green line. **(E)** The result ofimage segmentation based on ST_3_ segmentation of the GSM blue curve.

**Figure 2.**
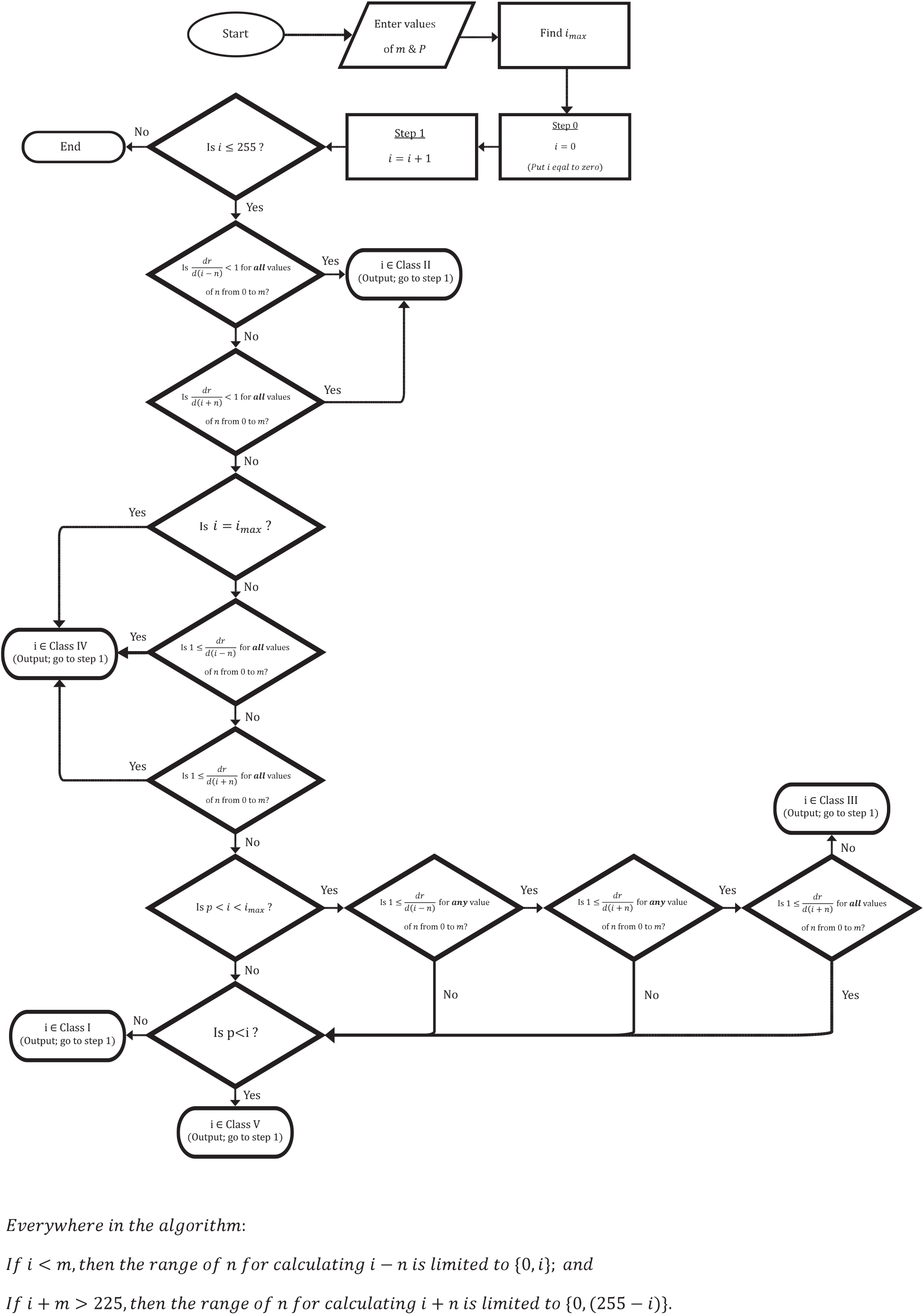
Flowchart of the classification algorithm developed for segmentation type 3. “*i*” represents the level of green color on the GSM graph (the variable shown on the horizontal axis). The variables *m*, *n*, and *p* belong to the set of natural numbers. *m* is the number of neighborhoods of *i* included in the process of determining the class of *i* and was set to 12; *p* is a threshold defined to distinguish the elements of the class I from the classes III and V and here was set to 50. *i_max_*: the level of *i* at which the amount of (*dr/di*) is maximum on the curve.

#### 2.3.4. Relationship between GSM attributes and canopy traits

In order to explore the potential relationships between various GSM-curve derived attributes and canopy traits, and also for evaluating capability of GSM in recognizing canopy status, performance of GSM in prediction of some quantitative and qualitative characteristics were tested using several data mining techniques including Decision trees (DT), Deep learning (DL), General linear model (GLM), Gradient boosted trees (GBT), Neural networks (NN), and Random Forest (RF; Mierswa *et al.*, 2006; analyses were carried out using RapidMiner Studio version 9.6). Since the field experiment was designed for other purposes and terminated before the image processing evaluations (see Haghshenas and Emam, 2019D), canopy characteristics would be defined and selected among previously specified measurements or subjections to environmental conditions. Accordingly, canopy coverage (CC), Normalized difference vegetation index (NDVI, measured only in the second season), canopy temperature, and grain yield were selected as quantitative traits, and subjecting to water and cold stresses, relative time of irrigation event, and also imaging date (Dataset) were defined as the qualitative traits for analyses.

Canopy coverage was calculated as below (Li *et al.*, 2010):

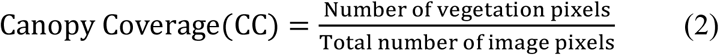

NDVI and canopy temperature were measured exactly before imaging using GreenSeeker (Trimble^®^, USA) and infrared thermometer (Terminator^®^ TIR 8861), respectively. Since repeated measuring of canopy temperature often lasted about 30 minutes, before using in the datamining analyses, readings of infrared thermometer were normalized by the method described by Haghshenas *et al.* (2019D), for minimizing the effect of fluctuations in environmental temperature. Moreover, as described before, 3 out of 6 replicates (45 plots) were exposed to deficit-irrigation. This treatment provided the opportunity of defining well- and deficit-irrigation as qualitative traits. Moreover, since imaging practices were scheduled to be regularly carried out (exactly) before and also (24 to 48 hours) after irrigation, time of imaging relative to the nearest irrigation events was defined as another qualitative trait. Occurrence of cold stress (belonging to Dataset D), and imaging date (recalling the correct dataset) were two other qualitative characteristics used for evaluating the performance of datamining models based on GSM attributes.

Ten-fold cross-validation (K=10) was used in Rapidminer to validate the results of datamining models. Moreover, default values and settings of the software were used for running the models. Input of datamining models included two different types of GSM-derived characteristics, based on which separate analyses were carried out:

a. *GSM-ST_2_ attributes*: the set of mean values make the red and blue curves of GSM graph. Depended on the degree of completeness of the curves, this set may include at most 510 values (255 points on each curve).
b. *Coefficients of GSM exponential equations (Exp. analysis)*: in this analysis, only the total number of 4 coefficients of the exponential equations fitted to the red and blue curves (*a*_*R*_, *a*_*B*_, *b*_*R*_, *b*_*B*_; see Equation 1) were used as the GSM-derived characteristics.

Statistical analyses were carried out using IBM SPSS Statistics for Windows (Version 19.0, Armonk, NY: IBM Corp.) and RapidMiner. Comparison and reconciliation between images and the ST_3_-derived graph segments were performed using Fiji and MATLAB, and some casual evaluations were carried out by XLSTAT (Version 2016.02.28451; Addinsoft). For keeping consistency among the figures of the present manuscript, everywhere an identical image and its GSM graph was used as example (which was captured from the experimental plot of the monoculture of the 1^st^ cultivar in the second replicate, Dataset A), unless mentioned otherwise.

## 3. Results

### 3.1. The overall form of GSM graph (ST_2_)

As shown in Fig. 1A & B, in the GSM graph of wheat canopy, the trends of mean red and mean blue values formed two smooth upward exponential shaped curves, among which the blue curve had a higher degree of curvature. Similar pattern could be observed for the images of all plots and datasets. The red and blue curves showed robust fit to exponential and polynomial equations. A comparison was carried out between the exponential and polynomial types of curve fittings for 24 images taken from the monoculture plots of Dataset A (data not shown), which revealed that the exponential trend of the blue curve had higher fitness compared to the red curve, while the fitness of polynomial equations of the both curves were almost similar. However, the coefficients of the exponential equations of both red and blue curves showed higher stabilities, compared with the coefficients of polynomial equations.

Supplementary Figure 1 (S1) indicates an interesting aspect of GSM curves. Irrespective of the dataset used, there was a strong negative linear correlation between the coefficients (*a* and *b*) of each exponential equation fitted to the red or blue curve. It is noteworthy that coefficients *a* and *b* determine the vertical intercept and curvature of exponential trend of the curve, respectively; so this observation revealed that as the y-intercept of the trend increases, its curvature decreases.

### Segmentation type 3 (ST_3_)

Evaluating the comparative local slopes (differentiations) of the red and blue trends relative to the 1:1 green line, 5 apparent parts (classes) on either red or blue curves were distinguished, based on which, consequently another type of segmentation was carried out (ST_3_, i.e. GSM curve-based segmentation, Fig. 1C). Also reconciling these curve-derived classes to the images by thresholding (Fig. 1C-E & Fig. S2) and visual assessments, five distinct corresponding segments were recognized on the green (vegetation) parts of the images, as described below (from green level of 255 towards 1):

- The highly reflective sun-exposed vegetation surfaces i.e. include the lightest pixels of the green canopy due to their spatial configurations relative to the sunlight and camera (based on bidirectional reflectance distribution function, BDRF). In this image segment, angles of the green surfaces are relatively homogeneous and show low variations. On the GSM curve, the red or blue trends in the range of this ST_3_ class, has non-stable local slopes greater than 1, which eventually at the left side extreme, approach the maximum declining rate (highest local slope) of the entire curve. This ST_3_ class is named “class V”, and the related curve range of the red and blue trends are shown as Red_45_ and Blue_45_, respectively.
- The image segment which covers all of the remained sun-exposed green surfaces with a complete set of altering angles. On this curve segment (i.e. class IV, with the curve range of Red_34_ /or Blue_34_), the sharpest declining slope of the GSM curve begins to decrease regularly towards the another threshold at which the local slope reaches that of the green line i.e. 1.
- The segment of partial shadow (class III, with the curve range of Red_23_ /or Blue_23_): Based on the visual assessments, this image segment seems to match the vegetation surfaces positioned under partial shadow (penumbra, Fig. 1 & S2). The GSM red and blue curves experience frequent fluctuations in relative local slopes around 1 (repeated slope shifting above and below the rate of the green line). It may imply that under partial shadow, the spectral behavior of vegetation in the red and blue bands are almost similar to the green one.
- The segment of complete shadow (class II, with the curve range of Red_12_ /or Blue_12_): this image segment includes the vegetation surfaces casted by shadow, i.e. lacks any sun-exposed point. Here, green color of vegetation surfaces is visually recognizable. This curve segment is the widest one across the GSM curve (includes the most diverse green values on the horizontal axis) with steadily declining relative slopes less than 1, i.e. has less reductions in the level of red and blue colors compared with green (this trend may also be comparable with the reported evidences for further penetration of green light into the leaf, as a particular micro-canopy; Sun *et al.*, 1998; Terashima *et al.*, 2009).
- The segment of deep shadow (class I, with the curve range of R_01_ or B_01_): this image segment is composed of the darkest green pixels of the canopy in the image. On the GSM curve, the local slopes of red and blue trends exceed again the constant slope of 1 (i.e. that of the green linear trend). Finally, intensity of blue light reaches a non-detectable reflection earlier than red on the curve, i.e. maybe less penetration to the deepest parts of the canopy.

Remarkably, the five ST_3_-derived classes described are complementary to each other, both in the image and on the curve; therefore, their integration cover all of the vegetation pixels in the image i.e. crop coverage. Moreover, while the separate implementation of the ST_3_ segmentation on the red and blue curves yielded a highly overlapped curve- and/or image-segments, yet, there were frequent finely differences between red and blue segments, which indicates that the shading pattern (and thresholds) is a band depended concept. This evidence, together with the precise differentiation of the canopy pixels in the class III and II (penumbra vs. complete shadow, Fig. S2), may indicate the relative efficacy of the ST_3_ segmentation in shading pattern evaluations. Supplementary Figure 3 shows some additional properties of the GSM graph, particularly with regard to ST_3_.

**Figure 3.**
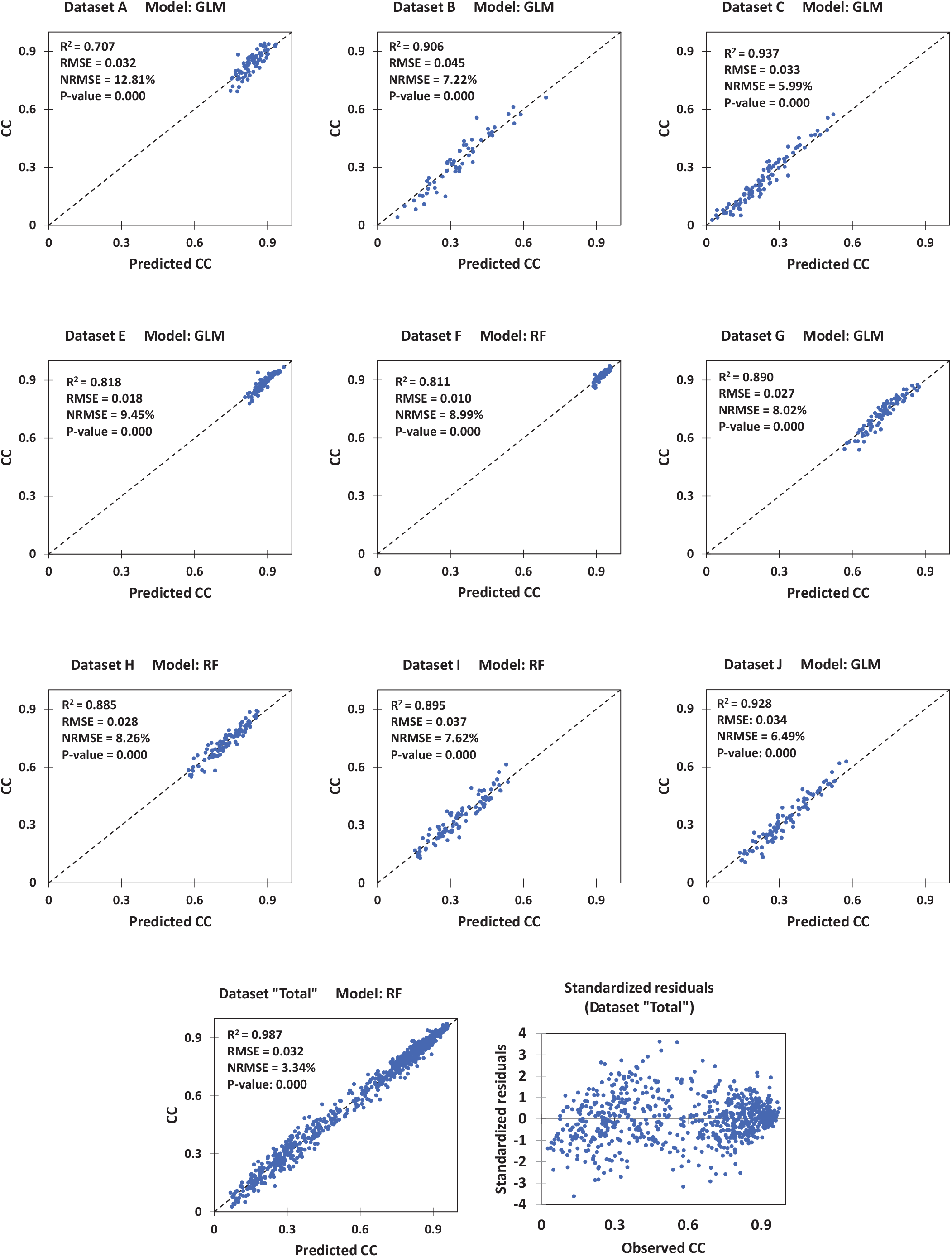
Cross-validation for prediction of canopy coverage (CC) by datamining models using GSM-ST2 attributes. The dataset “Total” is the combination of all other datasets. For each dataset, results of the model with the best performance are shown. GLM: General Linear Model; RF: Random Forest. NRMSE: normalized root mean square error (calculated by dividing the RMSE by the range of observed values).

### 3.3. Datamining analyses

#### 3.3.1. Based on GSM-ST_2_ attributes

In order to evaluate the capability of GSM graph properties to be used as HTP indices, their performance in prediction of some canopy traits were assessed using statistical and datamining approaches. As shown in Fig. 3, datamining models particularly RF and GLM predicted crop coverage precisely. The linear correlations between the observed and predicted CCs were very significant (P < 0.0001) irrespective the dataset (imaging date), and normalized root mean square errors (NRMSE) of prediction were mostly less than 10%, which decreased down to 3.34% in the analysis conducted using the combination of all datasets (i.e. Dataset “Total”).

Almost similar to CC, NDVI values were also predicted by RF and GLM models with high degrees of accuracy (Fig. 4). Besides the significant correlations between the predicted and observed values, NRMSE for predictions of single datasets ranged between 9.83% to 13.81% for Datasets I and F, respectively. When the combination of datasets (Dataset “Total”) was used, R^2^ raised to 0.96 and NRMSE fell to the minimum value of 5.19%. On the basis of RMSE, accuracy of NDVI prediction was not lower than 97.4% which resulted when Dataset G was used (in theory, NDVI has the range of 0 to 1).

**Figure 4.**
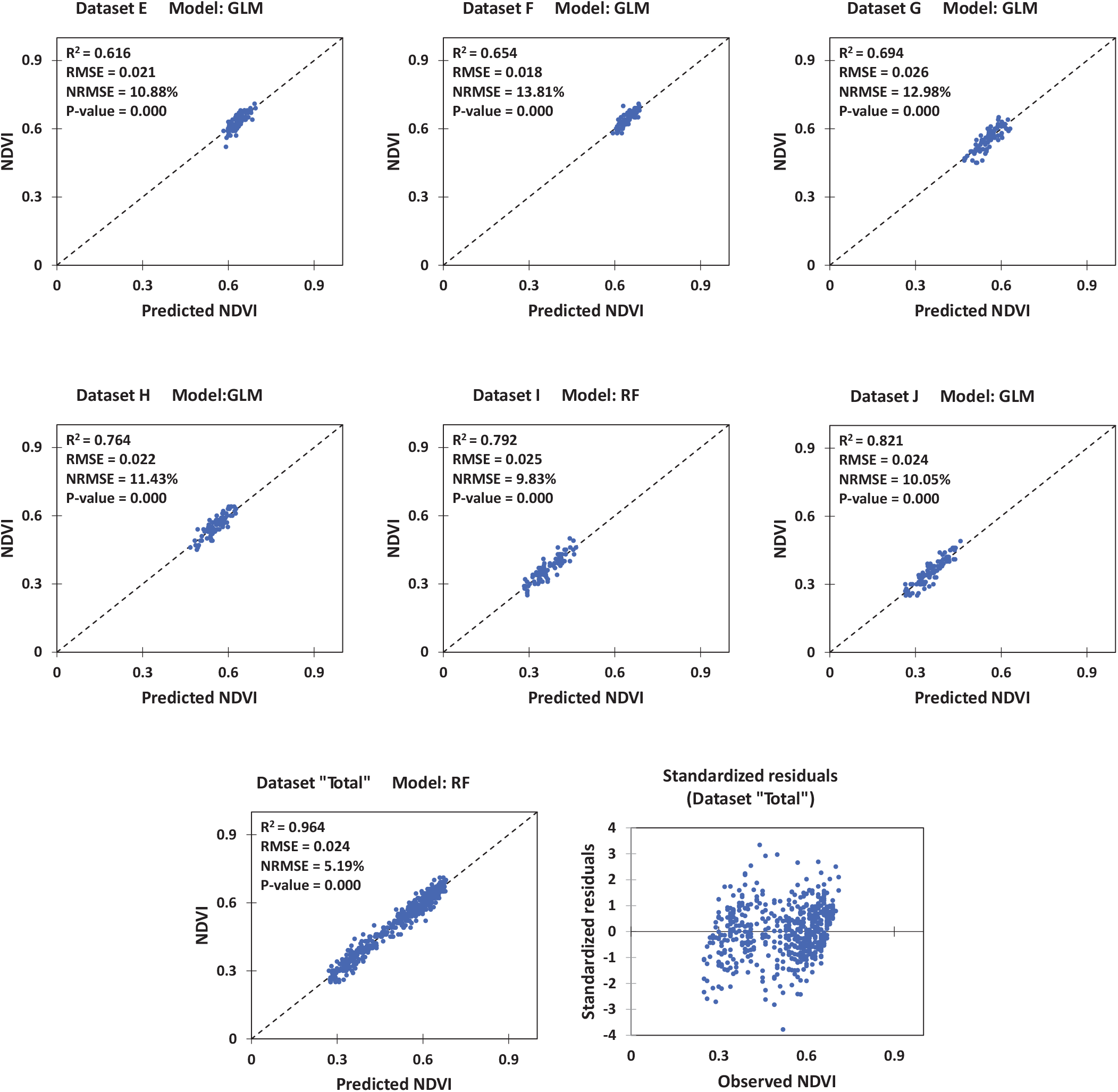
Cross-validation for prediction of Normalized Difference Vegetation Index (NDVI) by datamining models using GSM-ST2 attributes. The dataset “Total” is combination of all other datasets. For each dataset, results of the model with the best performance are shown. GLM: General Linear Model; RF: Random Forest. NRMSE: normalized root mean square error (calculated by dividing the RMSE by the range of observed values).

Although solely utilization of RGB images for estimating canopy temperature is not a common practice, and also the results of predictions based on the data of single datasets supported this fact (Fig. 5), correlations between the GSM-based predicted and observed temperatures were very significant in analyses of 6 out of 9 single datasets. However, when number of samples increased to 776 in Dataset Total, Random Forest model predicted canopy temperatures with a considerably higher accuracy (NRMSE = 8.01% & R^2^ = 0.872) or with error of less than 2 °C (RMSE = 1.89) for prediction of canopy temperature in a relatively broad range of 17.7 °C to 41.3 °C recorded from booting to the end of season.

**Figure 5.**
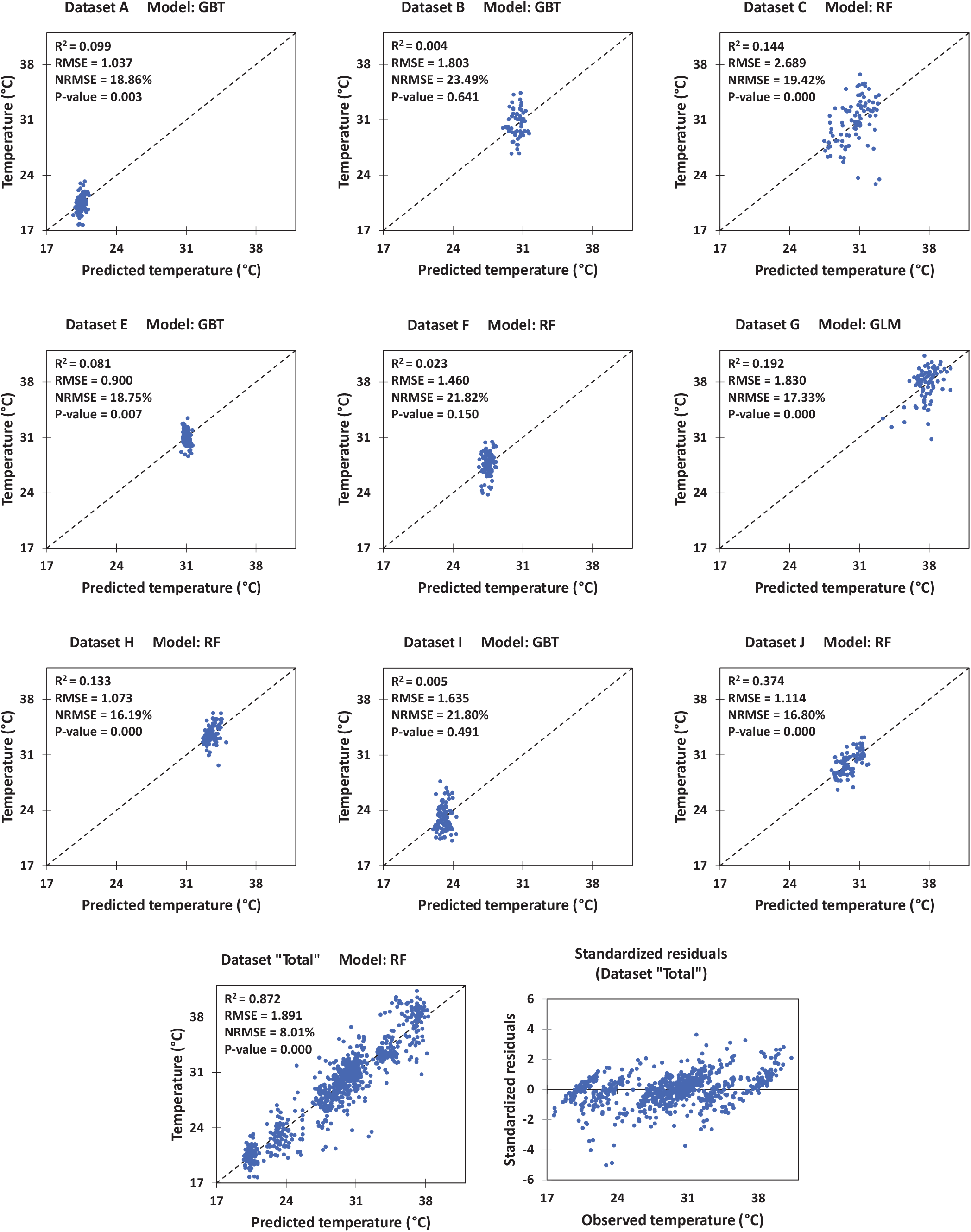
Cross-validation for prediction of canopy temperature (°C) by datamining models using GSM-ST2 attributes. The dataset “Total” is combination of all other datasets. For each dataset, results of the model with the best performance are shown. GBT: Gradient Boosted Trees; GLM: General Liniear Model; RF: Random Forest. NRMSE: normalized root mean square error (calculated by dividing the RMSE by the range of observed values).

Predictions of grain yield (GY) based on GSM-ST2 attributes were not satisfactory (Fig. 6). However, analyses of 6 out of 10 individual datasets showed significant correlations between the observed and predicted values. Moreover, when the last four datasets of the second year were used, results were progressed. This relative advantage of the late-season imaging dates was also somehow recognizable in the first year, as the last dataset (Dataset C) was the only one that showed significant correlation in this year (P-value = 0.009).

**Figure 6.**
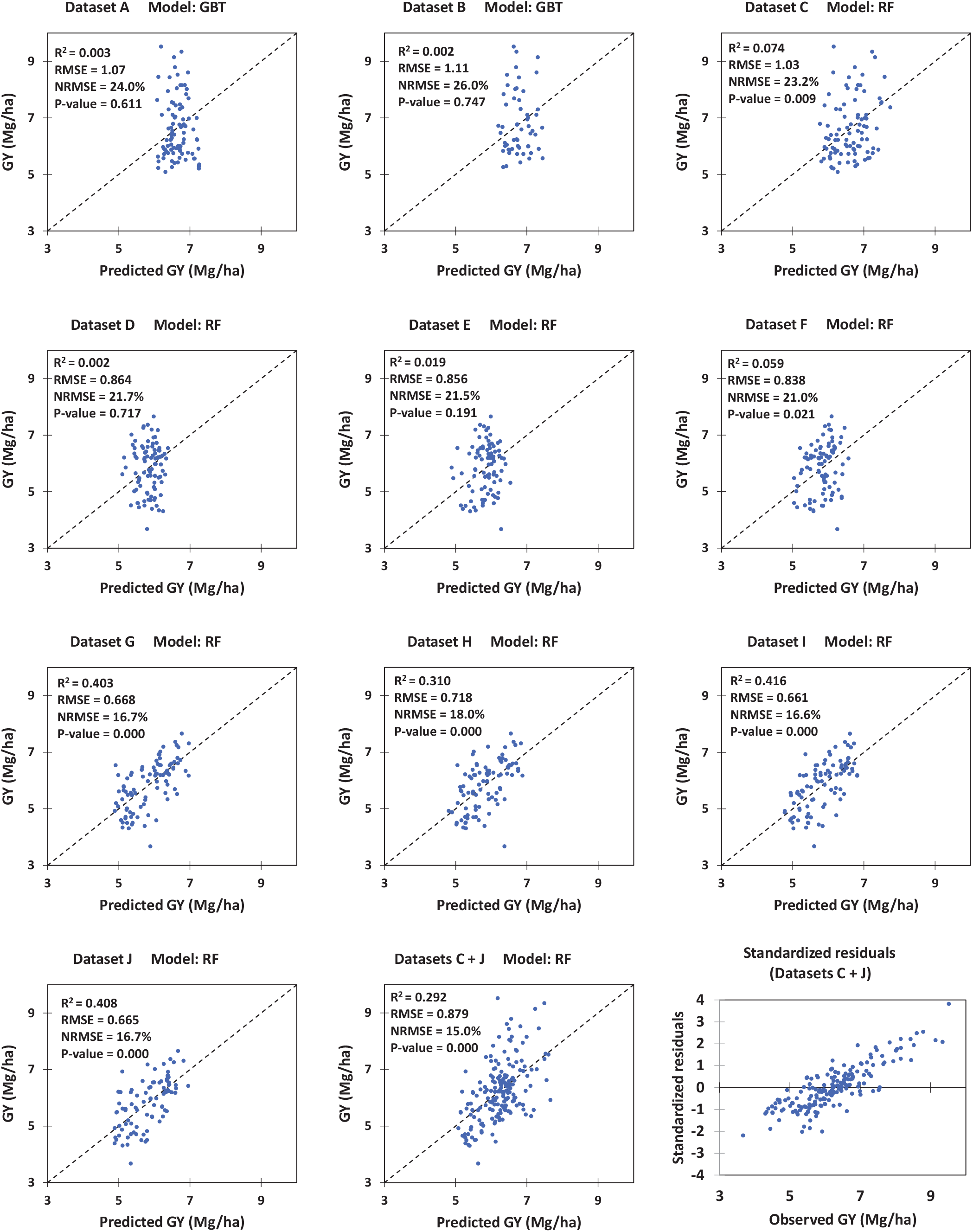
Cross-validation for prediction of grain yield (GY) by datamining models using GSM-ST_2_ attributes. As datasets C and J (included the images taken at the last imaging dates of the 1^st^ and 2^nd^ seasons) indicated the highest relationships in each season, their combination was also tested as a single dataset named “C+J”. For each dataset, results of the model with the best performance are shown. GBT: Gradient Boosted Trees; RF: Random Forest. NRMSE: normalized root mean square error (calculated by dividing the RMSE by the range of observed values).

Besides the analyses of quantitative traits of canopy, performance of GSM-ST_2_ attributes was also assessed in classification of the experimental plots based on the qualitative characteristics. As shown in Table 2, GBT and RF models classified the plots based on subjecting to water stress, cold stress, and time of irrigation, with 75.47%, 100%, and 89.08% accuracies, respectively. Moreover, in another analysis, the experimental plots were classified by RF model based on the date of imaging (dataset) with an overall accuracy of 92.84% (Table 3), which besides showing the GSM performance, may indicate the considerable effect of the interaction between crop phenology and illumination on images. Figure 7 shows the significant effects of these factors on ST_2_ attributes, which also may be implied as the comparative contribution of ST_2_ attributes in sensibility of GSM to the reported qualitative factors. Supplementary Figure 4 represents some example outputs (trees) of datamining models used for assessment of GSM performance in prediction of quantitative or qualitative characteristics of canopy.

**Table 2.** Performance of datamining models in classification of experimental plots based on qualitative traits using GSM-ST2 attributes.

| Comparison | Best model | Contrast groups |  | Model Performance |  |  |  |  |  |  |
| --- | --- | --- | --- | --- | --- | --- | --- | --- | --- | --- |
|  |  | I | II | Overall accuracy | Micro average accuracy | Prediction | Observation | True I (No. of plots) | True II (No. of plots) | Class precision |
| Water stress | GBT | WI | DI | 75.47%<br>+/- 4.54% | 75.49% | Pred. I (No. of plots) |  | 196 | 63 | 75.68% |
|  |  |  |  |  |  | Pred. II (No. of plots) |  | 61 | 186 | 75.30% |
|  |  |  |  |  |  | Class recall |  | 76.26% | 74.70% |  |
| Cold stress | RF | CS | U | 100% +/- 0% | 100% | Pred. I (No. of plots) |  | 90 | 0 | 100% |
|  |  |  |  |  |  | Pred. II (No. of plots) |  | 0 | 776 | 100% |
|  |  |  |  |  |  | Class recall |  | 100% | 100% |  |
| Irrigation event | GBT | Pre-Irr. | Post-Irr. | 89.08%<br>+/- 2.54% | 89.07% | Pred. I (No. of plots) |  | 281 | 30 | 90.35% |
|  |  |  |  |  |  | Pred. II (No. of plots) |  | 45 | 330 | 88% |
|  |  |  |  |  |  | Class recall |  | 86.20% | 91.67% |  |

The tested models include: Decision tree (DC), Deep learning (DL), General linear model (GLM), Gradient boosted trees (GBT), and Random forest (RF).
WI: well-irrigated; DI: Deficit-irrigated; CS: cold-stressed (plots); U: Unstressed (plots); Pre-Irr. : Pre-irrigation; Post-Irr. : Post-irrigation.

**Table 3.** Performance of Random Forest (the best model) in classification of experimental plots based on imaging dates (datasets) using GSM-ST_2_ attributes.

| Observation<br>Prediction | True A | True B | True C | True D | True E | True F | True G | True H | True I | True J | Class<br>precision<br>(%) |
| --- | --- | --- | --- | --- | --- | --- | --- | --- | --- | --- | --- |
| Pred. A | 90 | 0 | 0 | 0 | 0 | 0 | 0 | 0 | 0 | 0 | 100 |
| Pred. B | 0 | 29 | 6 | 0 | 0 | 0 | 0 | 0 | 0 | 1 | 80.56 |
| Pred. C | 0 | 19 | 83 | 0 | 0 | 0 | 0 | 0 | 1 | 3 | 78.3 |
| Pred. D | 0 | 0 | 0 | 90 | 0 | 0 | 0 | 0 | 0 | 0 | 100 |
| Pred. E | 0 | 0 | 0 | 0 | 84 | 5 | 0 | 1 | 0 | 0 | 93.33 |
| Pred. F | 0 | 0 | 0 | 0 | 6 | 85 | 0 | 0 | 0 | 0 | 93.41 |
| Pred. G | 0 | 0 | 0 | 0 | 0 | 0 | 84 | 5 | 0 | 0 | 94.38 |
| Pred. H | 0 | 0 | 0 | 0 | 0 | 0 | 6 | 84 | 0 | 0 | 93.33 |
| Pred. I | 0 | 2 | 1 | 0 | 0 | 0 | 0 | 0 | 89 | 0 | 96.74 |
| Pred. J | 0 | 6 | 0 | 0 | 0 | 0 | 0 | 0 | 0 | 86 | 93.48 |
| Class recall (%) | 100 | 51.79 | 92.22 | 100 | 93.33 | 94.44 | 93.33 | 93.33 | 98.89 | 95.56 | - |
| **Overall accuracy:** 92.84% +/- 2.91% |  |  |  |  |  | **Micro average accuracy:** 92.84% |  |  |  |  |  |

The values in the table are number of experimental plots.

**Figure 7.**
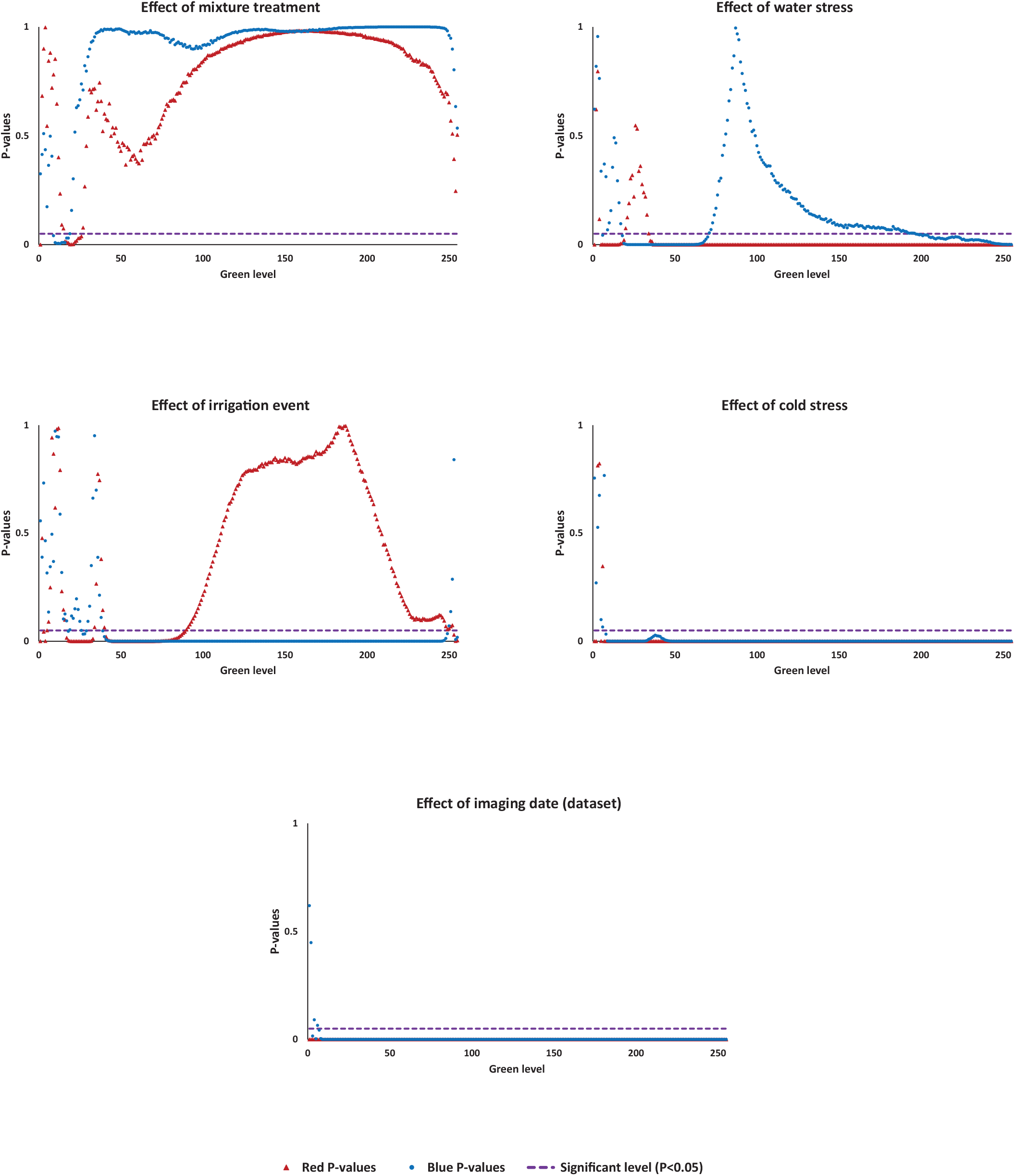
Significant effects of various qualitative factors (mixture treatment, water stress, irrigation event, cold stress, and imagin date) on GSM-ST2 attributes. The values below the dashed line show significant (P < 0.05) effects of the factors.

#### 3.3.2. Based on the coefficients of GSM exponential equations (Exp. analysis)

As a whole, performances of datamining models based on the coefficients of GSM exponential equations, followed the patterns reported in the previous part (i.e. GSM-ST_2_ based method); except that the accuracies declined slightly. In prediction of CC, GLM and RF models provided the best predictions with the NRMSE values ranges between 6.96% to 14.72% for individual datasets, and the minimum value of 4.44% when the combination of all datasets was used (Fig. S5). Very significant correlations were also observed between the predicted and measured NDVI values irrespective of the dataset; however, prediction accuracy increased toward the end of season. Similar to the results of GSM-ST_2_ method, the highest accuracy was achieved when the combination of all datasets was used (NRMSE= 6%). Although the correlation between the observed and predicted canopy temperatures were significant in five out of the nine datasets evaluated, the prediction accuracy was relatively acceptable only in the combined dataset where the error declined to 2.29 °C (NRMSE = 9.72%, R^2^ = 0.811; Fig. S7). As shown in Fig. S8, except the datasets included the images taken before spike emergence in two seasons (i.e. Datasets A & D), the correlation between the observed and predicted grain yields were significant. However, the error of prediction never fell below 0.667 kg/ha in the individual datasets (NRMSE ≥16.73%). Performance of datamining models in *Exp. analyses* may also be explained by the data represented in Table 4, which shows the correlations among the canopy quantitative traits and coefficients of GSM exponential equations. Most of correlations were very significant (P-value < 0.001), and the lower ones were observed in GY prediction. Correlations of CC or NDVI with coefficients of the red trend were apparently higher compared with the blue trend, which in contrast, indicated a comparatively higher relationship with canopy temperature. Accordingly, curvature of red trend (*b*_*R*_) indicated strong positive correlations with CC and NDVI, but a negative and comparatively lower relationship with canopy temperature.

**Table 4.** Correlations (R values) among the quantitative traits of canopy and coefficients of GSM exponential equations.

| Factors | $a_{Red}$ | $b_{Red}$ | $a_{Blue}$ | $b_{Blue}$ |
| --- | --- | --- | --- | --- |
| CC | **-0.927** | **0.871** | **0.240** | **-0.176** |
| Temp. | **0.220** | **-0.172** | **0.455** | **-0.402** |
| NDVI | **-0.926** | **0.859** | **0.474** | **-0.420** |
| GY | -0.165 * | 0.107 ns | 0.204 * | **-0.306** |

Correlations are carried out using the corresponding datasets entitled "Total" (for CC, temperature, and NDVI) and C+J (for grain yield). CC: canopy coverage; Temp. : canopy temperature; NDVI: Normalized difference vegetation index; GY: grain yield. $a_{Red}$ , $b_{Red}$ , $a_{blue}$ , and $b_{blue}$ are the coefficients of GSM exponential equations (see manuscript).
\* significant correlation (p-value <0.05); ns: not significant; and values in bold indicate very significant correlations (p-value <0.001).

Accuracies of classification of experimental plots based on subjecting to water stress, cold stress, and time of irrigation were 64.95%, 100%, and 73.78% respectively (Table 5) which shows 10.52%, 0%, and 15.3% reductions, respectively compared with the results of GSM- ST_2_ analysis. Similarly, the overall accuracy of classification based on the imaging date (dataset) declined to 80.95% (Table 6). Table 7 indicates the significant effects of qualitative factors (treatments, stresses, and imaging date) on the coefficients of GSM exponential equations. Mixture treatment and water stress had not any significant effect on the coefficients, while the effects of other factors were very significant (P < 0.001). These results may reveal the reason of comparatively higher degrees of accuracy in classifications based on the cold stress, time of irrigation, and imaging date. Supplementary Figure 9 represents example outputs (trees) of datamining models used in the *Exp*. analyses.

**Table 5.** Performance of datamining models in classification of experimental plots based on qualitative traits using coefficients of GSM exponential equations.

| Comparison | Best model | Contrast groups |  | Model Performance |  |  |  |  |  |
| --- | --- | --- | --- | --- | --- | --- | --- | --- | --- |
|  |  | I | II | Overall accuracy | Micro average accuracy | Prediction \ Observation | True I (No. of plots) | True II (No. of plots) | Class precision |
| Water stress | GBT | WI | DI | 64.95% +/- 5.37% | 64.95% | Pred. I (No. of plots) | 272 | 152 | 64.15% |
|  |  |  |  |  |  | Pred. II (No. of plots) | 120 | 232 | 65.91% |
|  |  |  |  |  |  | Class recall | 69.39% | 60.42% | - |
| Cold stress | GLM (and DL) | CS | U | 100.00% +/- 0.00% | 100% | Pred. I (No. of plots) | 90 | 0 | 100% |
|  |  |  |  |  |  | Pred. II (No. of plots) | 0 | 776 | 100% |
|  |  |  |  |  |  | Class recall | 100% | 100% | - |
| Irrigation event | GBT | Pre-Irr. | Post-Irr. | 73.88% +/- 6.80% | 74% | Pred. I (No. of plots) | 243 | 96 | 71.68% |
|  |  |  |  |  |  | Pred. II (No. of plots) | 83 | 264 | 76% |
|  |  |  |  |  |  | Class recall | 0.745 | 0.733 | - |

The tested models include: Decision tree (DC), Deep learning (DL), General linear model (GLM), Gradient boosted trees (GBT), and Random forest (RF).
WI: well-irrigated; DI: Deficit-irrigated; CS: cold-stressed (plots); U: Unstressed (plots); Pre-Irr.: Pre-irrigation; Post-Irr.: Post-irrigation.

**Table 6.** Performance of General Linear Model (the best model) in classification of experimental plots based on imaging dates (datasets) using coefficients of GSM exponential equations.

| Observation<br>Prediction | True A | True B | True C | True D | True E | True F | True G | True H | True I | True J | Class<br>precision<br>(%) |
| --- | --- | --- | --- | --- | --- | --- | --- | --- | --- | --- | --- |
| Pred. A | 90 | 0 | 0 | 0 | 0 | 0 | 0 | 0 | 0 | 0 | 100% |
| Pred. B | 0 | 29 | 9 | 0 | 0 | 0 | 0 | 0 | 0 | 0 | 76.32% |
| Pred. C | 0 | 23 | 65 | 0 | 0 | 0 | 0 | 0 | 0 | 15 | 63.11% |
| Pred. D | 0 | 0 | 0 | 90 | 0 | 0 | 0 | 0 | 0 | 0 | 100% |
| Pred. E | 0 | 0 | 0 | 0 | 76 | 14 | 0 | 0 | 0 | 0 | 84.44% |
| Pred. F | 0 | 0 | 0 | 0 | 10 | 76 | 0 | 0 | 0 | 0 | 88.37% |
| Pred. G | 0 | 0 | 0 | 0 | 0 | 0 | 57 | 37 | 0 | 0 | 60.64% |
| Pred. H | 0 | 0 | 0 | 0 | 4 | 0 | 32 | 53 | 0 | 0 | 59.55% |
| Pred. I | 0 | 3 | 2 | 0 | 0 | 0 | 1 | 0 | 90 | 0 | 93.75% |
| Pred. J | 0 | 1 | 14 | 0 | 0 | 0 | 0 | 0 | 0 | 75 | 83.33% |
| Class recall (%) | 100% | 51.79% | 72.22% | 100% | 84.44% | 84.44% | 63.33% | 58.89% | 100% | 83.33% | - |
| **Overall accuracy: 80.95% +/- 3.50%** |  |  |  |  |  | **Micro average accuracy: 80.95%** |  |  |  |  |  |

The values in the table are number of experimental plots.

**Table 7.** Significant effects of qualitative factors on coefficients of GSM exponential equations.

| Factor (treatment) | P-value |  |  |  |
| --- | --- | --- | --- | --- |
| | $a_{Red}$ | $b_{Red}$ | $a_{Blue}$ | $b_{Blue}$ |
| Mixture | 0.928 | 0.938 | 0.966 | 0.856 |
| Water stress | 0.055 | 0.093 | 0.942 | 0.281 |
| Irrigation event | 0.131 | **0.001** | **0.000** | **0.000** |
| Cold stress | **0.000** | **0.000** | **0.000** | **0.000** |
| Imaging date (dataset) | **0.000** | **0.000** | **0.000** | **0.000** |

### 3.4. Effect of Phenology on the shape of GSM curves

Figures 8 A, S10, and S11 show the variations of GSM curves during the growing season. Accordingly, from emergence, the overall areas between green-red and green-blue curves begin to increase up to their maximum values, with the most curvature form at the late tillering and early stem elongation phases, at the time when the most dense canopy has been formed due to intensive leaf production and low plant heights. Following stem elongation, increased canopy height lead to a returning trend, as towards the end of growing season -in particular- the red trend line again approaches to the green line. This evidence indicates the potential capability of GSM graph in monitoring and quantifying the phenological trends, particularly considering the sensibilities kept even late in the season.

**Figure 8.**
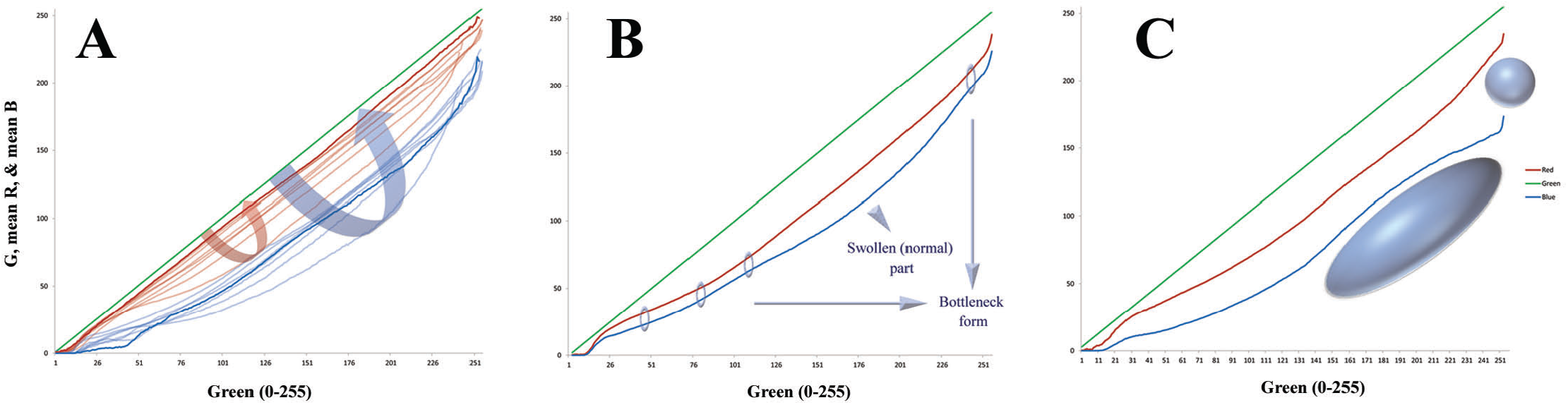
Changes of the GSM graph as influenced by phenology and environmental stresses. The red and blue curves are affected by **(A)** canopy growth stage (phenology^)^, **(B)** water and **(C)** cold stresses, respectively. whose diversion from the normal forms are indicated for the latter two as a bottleneck form or by ellipsoids. For more details, see Supplementary Figures 10 to 13.

### 3.5. Effects of water and cold stresses on the shape of GSM curves

As shown in Fig. 8 B&C, Fig. S12, and Fig. S13, it was also observed that the shape of GSM graph was influenced by environmental stresses i.e. comparatively abnormal diverted forms mainly made by altered (either increased or decreased) reflection of blue light under full sunlight to complete shadow conditions. Such observations may propose an acceptable diagnostic role for GSM in detecting various kinds of stresses which affect canopy optics or BDRF, especially considering the total or partial variations of the area between the blue-red curves and blue-green trends. According to the findings, it should be emphasized that ST_3_ is developed based on the images of wheat canopies grown under relatively optimal conditions, and thus it seems that other types of curve-segmentation may be required for stressful conditions.

## 4. Discussion

In the present study, the capability of a simple 2D image mining model in characterizing the shading pattern of crop canopies, and also providing a source of vegetation indices for HTP was evaluated; which is based on the canopy segmentation according to the linear gradient of green color. While the 3D-based methods of HTP require multi-directional imaging and sophisticated computation algorithms (see Artzet *et al.*, 2019; Li *et al.*, 2019; Paulus *et al.*, 2019; and Shi *et al.*, 2019), quantitative analyzing of the shading pattern recorded in a single 2D image has the potential to provide indirectly valuable data of canopy; even though, the conventional 3D morphological characteristics are not considered or estimated directly. Various analyses conducted here, indicated that GSM provides a simple solution both for characterizing the shading pattern and also for introducing a new pool of image-derived vegetation indices for crop HTP. The archive of images used were taken from the canopies of wheat cultivar mixtures, which could make the assessments more challenging, due to including higher intra-species heterogeneities compared with the monocultures (e.g. see Borg *et al.*, 2018, and Reiss & Drinkwater, 2018). However, it is indicated that the GSM model has acceptable performances in quantitative illustration of the reflection status above the canopy in a simple form, discrimination of canopy green surfaces according to their exposure to sunlight, e.g. distinguishing between partial- and complete shadows, and eventually prediction of various quantitative and qualitative properties of canopy with acceptable degrees of accuracies.

The GSM graph has an intrinsic simplicity arising from its simple structure, non-complex computation (which is based on a routine pixel classification and basic statistics), and straightforward interpretation i.e. according to the comparative variations in the local curve slopes, which may show the relative variations in RGB triplets under changing illumination. Despite such simplicities, GSM graph extracts and represents an orderly pattern from a naturally complex and disordered phenomenon i.e. the extremely sophisticated light scattering above the canopy. Based on this fact, GSM seems to go beyond a technique, and may be readily used as an identification criterion for vegetation canopies. In other words, in an ideal scope and after justified by adequate studies, a vegetation canopy may be quantitatively defined and distinguished from others, using the overall form of its GSM graph or equations of the curves. As described before, GSM graph is made based on a comprehensive collection of reflections from surfaces usually arranged and/or located in canopy in a sophisticated manner, thus exposed to the various degrees of sunlight. That is to say, assuming the whole canopy as a green flat homogenous surface unit, e.g. as a conceptual pixel, the GSM graph provides a comprehensive illustration of the visible spectral behavior of this unit from a fixed point of view, while it is experiencing a complete set of changes in illumination (quantified here as the range of RGB green level from 1 to 255). However, it should be noted that depended on the study purposes, sometimes canopy cannot be assumed as a single unit with homogenous reflectance behavior; e.g. due to the heterogeneities in the pattern of leaf senescence, and/or nitrogen distribution across the canopy height or between the sun-exposed vs. shaded surfaces (see Dreccer *et al.*, 2000; Bertheloot *et al.*, 2008; Hikosaka *et al.*, 2016; Kitao *et al.*, 2018). Under such conditions, the purpose of the study determines whether these variations should be neglected or taken into account. Anyway, the GSM graph indicates the variations in the relative reflections of the red and blue bands, regardless of the necessity for knowing the biophysiological background.

Variant illumination is often known as a problematic phenomenon in the image-based canopy studies; in particular, for segmentation of the vegetation parts from background (see Guo *et al.*, 2013; Yi *et al.*, 2015; and Chopin *et al.*, 2018). However, it is the foundation and main topic of the GSM approach. To the best of our knowledge, it is the first time that illumination is defined and evaluated as the gradient of a RGB triplet i.e. the linear trend of green levels, instead of using grayscale or other synthetic criteria. The idea was that, besides its simplicity, green color is the most rigorous reflection from vegetation, so it could distinguish among various illuminations more efficiently. In addition to the properties described for the overall form of GSM graph, a further analysis on the curves led to a curve-based segmentation approach (ST_3_), based on which the canopy was segmented into 5 classes from full-sunlight to deep-shadow, with acceptable accuracies (see Fig. S1). The criterion used for ST_3_ segmentation was consistent changes in local slopes of red or blue curves, relative to the fixed slope of the green trend i.e. equals 1. Accordingly, separate shading patterns and series of thresholding values were recognized for each of the red or blue curves; which implies that ST_3_ distinguishes between the red-color-based and blue-color-based shading, while their results are highly overlapped in most cases. However, it should be kept in mind that the equations and/or equation fitness of GSM curves may be changed under stressful and abnormal conditions. So the ST_3_ may also be required to be carried out in a different way using other purpose-oriented algorithms; e.g. through segmenting the GSM curves into only two segments based upon having slopes higher or lower than 1. Indeed, it seems that the GSM approach is flexible enough to provide the opportunity for calculating such diversions from the typical form.

One of the most important functions of GSM model was prediction of various quantitative and qualitative characteristics of canopy. Based on the type of GSM output (which was used as the input of datamining models), two parallel datamining approaches were evaluated i.e. using GSM-ST_2_ attributes and coefficients of the exponential equations fitted to the GSM curves (Exp. analysis). Although the first approach supported by up to 510 attributes had a comparatively better performance in the prediction and classification analyses, the second method also provided acceptable results particularly in the cases where the GSM-ST_2_ approach was successful too. Considering only 4 variables were used as the input of datamining models in the Exp. analysis, it may be remained as a considerable choice where the cost of time and/or computation is important (e.g. for developing smartphone apps). Besides, the strong negative correlation between the two coefficients of each curve may potentially allow even more reduction in the number of Exp. derived inputs needed for datamining models.

It is notable that the main purpose of conducting datamining analyses in the present study was validating the GSM model, and not necessarily introducing image-based alternatives for measuring the quantitative characteristics such as NDVI, canopy temperature, or CC, for which there are currently well-known and simple methods available. However, while inexpensive and precise devices such as infrared thermometer or GreenSeeker® are commonly used for measuring canopy temperature and NDVI, the possibility of replacing them in an integrative estimation approach based on processing a single image may also worth to be considered, e.g. for improving the efficiency of phenotyping systems by reducing number of sensors, or extracting various types of data from the images shared by smallholder growers.

Canopy coverage was predicted by GSM with high accuracies irrespective the datamining approach and dataset used. In spite of the fact that CC is a simple image-derived index with a distinct formula, it was also used for validation of GSM; because based on definitions, GSM is theoretically independent from canopy coverage and/or number of vegetation pixels. In other words, only several hundred green pixels taken from a small slice of the image may be enough to form the GSM graph. Nevertheless, high correlations between the actual and predicted CC suggested that GSM graph may be affected considerably by canopy coverage. As an evidence for this, correlations between CC and coefficients of the exponential trends (in Exp. analysis) were very significant and in particular, it had a strong positive (non-linear) relationship with curvature of the red trend (Table 4).

Acceptable performance of GSM in predicting NDVI and canopy temperature might be an unexpected result; since their measurements are partially to thoroughly independent from the reflection in visible band. However, high degrees of accuracy observed in prediction of NDVI, and acceptable levels of error obtained for prediction of canopy temperature in the case of using the combination of all datasets, suggested that GSM may even be utilized for conducting such unusual estimations. It seems that the reason for this capability lies in the potential effect of temperature dynamics on the spectral behavior of canopy in the range of visible light. Indeed, it is expected that GSM graph is affected indirectly by variations of temperature. Furthermore, a similar argument may be valid for the relationship between GSM-derived attributes and NDVI; except that besides the near-infrared portion, red color level also contributes equally to the NDVI calculation, which can directly improve the GSM sensibility. Utilizing genetic algorithms, Costa *et al.* (2020) developed an RGB-based index (i.e. *vNDVI* or visible NDVI) for estimating NDVI using airborne images, and reported an average error of 6.89% for predictions. To the best of our knowledge, this is the only study in the literature that has reported NDVI estimations only based on visible (RGB) data. Besides, no report was found for estimating canopy temperature merely using RGB cameras.

In general, predictions of GY were not satisfactory in the present study. In comparison with other quantitative properties evaluated here (i.e. CC, NDVI, and canopy temperature), GY is a more complicated trait which is determined in long term. Indeed, while the time scales of variations and measurement intervals for CC, NDVI, or canopy temperature may be from several minutes to days, GY is the result of the continues interaction between genotype and various environmental factors throughout the season. Therefore, it is not so unlikely that image-derived data has higher correlations with instantaneously measured traits, compared with GY. However, the very significant relationship observed between the actual and predicted values of GY in most of analyses, may be a promising sign that GSM-derived attributes are sensible to the mechanisms contribute to grain yield.

Among the qualitative characteristics of canopies, imaging date and a casual occurrence of cold stress were predicted with the highest degrees of accuracy, irrespective the type of GSM derived attributes. Accuracies of classification of the experimental plots based on exposing to water stress and time of irrigation were also considerable (Tables 2 & 5). These results may provide the opportunity for utilizing GSM for studying the short- or long-term effects of water stress and scheduling irrigation events based on RGB images. A notable point about sensitivity of GSM to imaging date is that most of the datasets used here were captured in various phenological stages during the post-anthesis period; so further studies are required to determine and separate the effects of illumination and phenology on GSM graph. Overall, results of datamining analyses support the idea that despite the simple structure and calculations, GSM has a multi- or general-purpose capability to cover a relatively broad spectrum of requirements in phenotyping studies.

Researches with the aim of evaluating the structure and optics of vegetation canopies, have utilized a relatively broad spectrum of physical bases, mathematical models, and practically, novel photometric instruments (e.g. Verhoef, 1984; Welles and Cohen, 1996; Gower *et al.*, 1999; Jonckheere I. *et al.*, 2004; Omasa *et al.*, 2006; Jacquemoud *et al.*, 2009; Zhao *et al.*, 2011; Glatthorn and Beckschäfer, 2014; Bauer *et al.*, 2016; Yao *et al.*, 2016). Despite the diversity and evolution of the methodologies, the two early introduced concepts of leaf area index –LAI-(Watson, 1947), and the exponential trend of light attenuation presented based upon the Beer-Lambert law (Monsi and Saeki, 2005), solely or in combination, have been the cornerstones of almost every conventional analyses in this context. Accordingly, leaves with their individual characteristics e.g. type, angle, and configuration, act as the light attenuating materials; and LAI is comparable with the material concentration. Conventional measurement approaches include estimating the conceptual index of LAI either by destructive sampling or indirect methods (Weiss *et al.*, 2004; Behera *et al.*, 2010; Liu *et al.*, 2010; Viña *et al.*, 2011; Mu *et al.*, 2017), and/or modeling the light extinction trend based on recorded light intensities by *in situ* sensors (Jonckheere *et al.*, 2004; Munier-Jolain *et al.*, 2013; Xue *et al.*, 2015; Perot *et al.*, 2017). For instance, in the well-established method of digital hemispherical photography or DHP, LAI is estimated based on the gap fraction of the images often taken using fisheye lenses (van Gardingen *et al.*, 1999; Jonckheere *et al.*, 2004; Kobayashi *et al.*, 2013; Zhao *et al.*, 2019). Of course, this approach is developing technically both in algorithms (see Kobayashi *et al.*, 2013; and Loffredo *et al.*, 2016) and imaging methods and tools; e.g. Confalonieri *et al.* (2013) introduced the first smartphone app for estimating LAI based on gap fraction. This application uses the smartphone sensors i.e. camera and accelerometer to automatically acquire images at 57.5° below the canopy while the user is rotating the device along its main axes. However, even in these methods, the results of remote sensing analyses are calibrated based on LAI or outputs of in-field commercial sensors, which may be a laborious practice or require utilization of complex models. In contrast, as practiced in the present study, the GSM graph may be interpreted independently of LAI and associated models, or additional sensors.

Although here the focus was basically on evaluating GSM as a 2D image mining approach for HTP, it seems to have the potential to contribute to canopy optics and biophysical assessments. Indeed, GSM is expected to have strong contribution with canopy BDRF and may be used for measuring the canopy radiation field and quantifying the scattering process inside vegetation canopies, which also requires to be compared with the results of valid physical methodologies in next studies based on the *in-situ* measurements, and/or combining the GSM output with developed radiative transfer models (e.g. see Vilfan *et al.*, 2016; Yang *et al.*, 2017; and Li *et al.*, 2018). It is notable that here, no in-field measurement was carried out for analyzing the canopy architecture properties, e.g. measuring PAR, or leaf area for LAI–estimations; since the idea of the study was formed after field harvest, and hence, the focus was basically put on evaluating GSM as a HTP platform for datamining analyses, for which the data was available to validate the GSM performance.

Green-gradient based segmentation model may provide new implications and definitions for characterizing vegetation canopies. For instance, canopy depth and/or layer may be defined using GSM as an optical term (e.g. based on GSM ST_3_ classes, or the green level itself) instead of being a spatial concept; because green surfaces may be completely subjected to either sunlight or shadow at any height in the canopy, though, the frequency or probability of exposures may be predictable based on the spatial height. Accordingly, depending on objectives, once segmented from background -i.e. after ST_1_-even a rolled leaf that casts a shadow on itself can be considered as an individual canopy with its own GSM graph, which also may be comparable with a wheat canopy or even with a forest in a satellite image. Although, even if such canopies of different tempo-spatial scales have similar GSM graphs or curve equations, their GSM graph should be individually interpreted with respect to the biophysical facts of the corresponding scale; so it may need additional conversions. Moreover, since the GSM approach is based on variability of illumination, it is recommendable for future researches to study the effects of associated factors such as gamma correction, sensor type, imaging condition, time, etc.

In summary, it is shown that the GSM model which is based on a simple image segmentation, has the potential to be utilized as a multipurpose HTP platform and canopy studies including evaluation of shading pattern, estimating physiological properties, and stress analyses. Also it seems to be capable to contribute to the evaluations of canopy optics with various degrees of dependence to the conventional methodologies and biophysical levels, e.g. in potential combination with radiative transfer models, or even as an independent approach for monitoring and assessment of image-derived properties. Yet, further studies are needed to improve our understanding of GSM and its various applications.

## 5. Conclusion

Considering the high overlap between the ranges of PAR and visible spectra, the low-cost and easily taken 2D ground-based nadir images of crop canopies have become the pivot elements of HTP approaches. In the present study the shading patterns of wheat canopies were studied using segmentation of canopy parts based on the gradient of green level of vegetation pixels in RGB color system. The sets of mean red and mean blue values in each green level formed two exponential upward curves, which along with the 1:1 green line –or gradient-were jointly introduced as the GSM graph. Based on the major shifts in the local slopes, five distinct parts were recognized on each curve. Segmentation of the GSM curves into these parts –i.e. ST_3_- and reconciling them to the images could effectively classify the pixels of crop coverage based on their exposure to sunlight. Moreover, using datamining approaches, it was indicated that GSM model has a considerable potential to predict various quantitative properties of wheat canopies (including CC, NDVI, and somewhat canopy temperature) with high degrees of accuracy. The model was also capable to classify experimental plots precisely based on exposure to cold and water stresses, imaging date, and irrigation time. Accordingly, effects of phenology and environmental stresses on the shape of GSM curves are discussed.

In conclusion, GSM is introduced as a multipurpose image mining platform with the potential utilization in:

- Quantitative representation of crop canopies in a simple form, e.g. as one or two equations, or even by the entire GSM graph;
- Recognition and classification of shading pattern inside the canopy;
- Prediction of various quantitative or qualitative characteristics of crop canopies using datamining models, with the potential application in HTP and screening programs;
- Assessing dynamics of canopy reflectance in the range of visible band, as affected by phenology, environmental stresses, or temperature.
- Studying canopy optics, e.g. for evaluation of the radiation field above the canopy, BDRF, and pattern of visible light scattering inside the canopy (only after validation by appropriate biophysical measurements or models).

It seems that the findings reported here, including the novel technique and concepts as well as the suggested applications, may improve our knowledge of crop canopy status and provide new vegetation indices for crop phenotyping.

## Acknowledgements

The authors wish to thank Shiraz University for providing field experiment facilities. Mr. Saeid Jafarizadeh is gratefully acknowledged for his contribution in writing MATLAB codes.

---

**Supplementary Figure 1.**
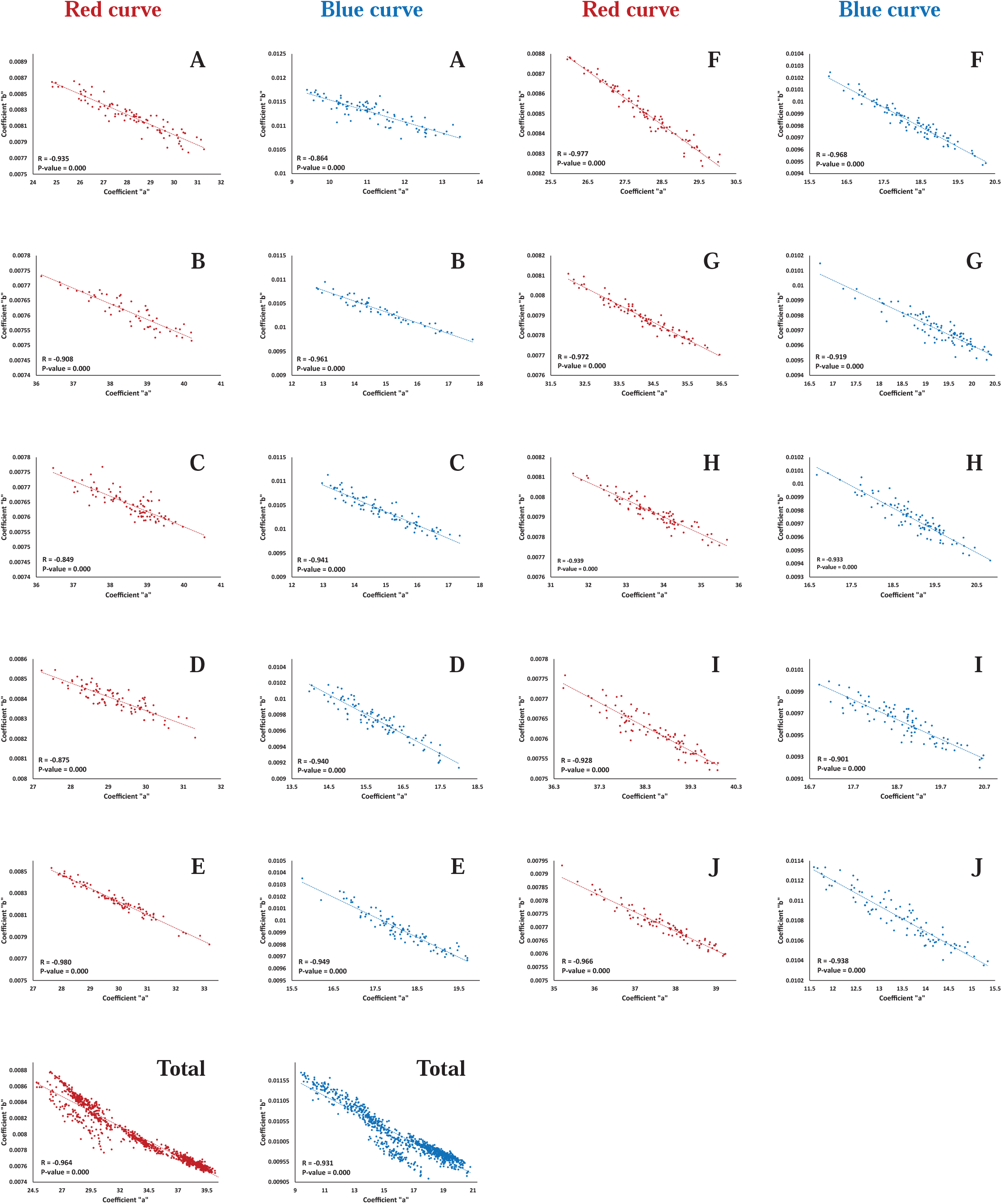
Negative linear correlations between the “*a*” and “*b*” coefficients of the exponential equations fitted to the GSM red- and blue-curves, respectively. *A* to *J* show the image datasets, and the dataset “Total” is the combination of all other datasets. It is notable that “*a”* and “*b”* determine the vertical intercept and curvature of each curve, respectively.

**Supplementary Figure 2.**
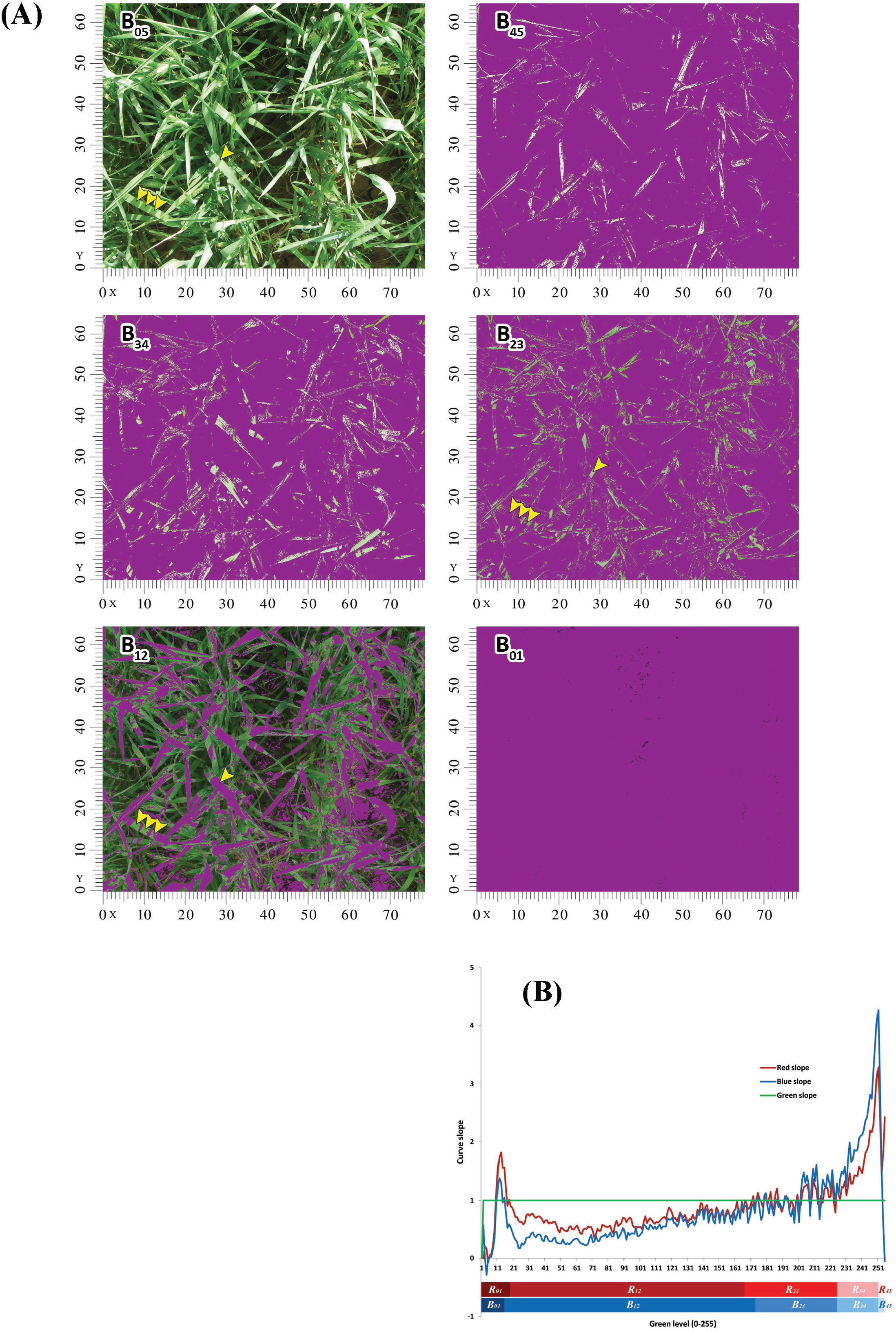
Segmentation type 3 (ST_3_) for the image ofa wheat canopy (monoculture ofthe 1^st^ cultivar) at booting stage. The central section ofan image segmented based on ST**_3_** ofthe blue curve (depended on variations in the curve local slopes relative to the green trend). In each figure, the purple mask has covered the excluded pixels. Compare the shadow on the leaf in the position (x=28, y=26) with the three parallel strips ofshadows on the leafat (x= 10, y=15). The first one is categorized as a partial shadow, since it is observable in the segment B23 (where the local slopes ofthe blue curve are almost equal to the slope ofgreen trend, i.e. 1), while, the second set is recognized as complete shadow (B12, i.e. the local slopes ofthe blue curve are lower than 1). Similarly, different parts of any single rolled or folded leaf in various segments may be tracked. Notably, (i) non-vegetative pixels are excluded previously by the segmentation type 1 (ST1), and (ii) the 5 segments (classes) demonstrated here are complementary to each other, so each pixel may be included only in one segment. **(B)** The relative variations in local slopes ofthe red and blue GSM curves, compared to the green line. ST_3_ is developed based on this graph; see Fig. 2 for details.

**Supplementary Figure 3.**
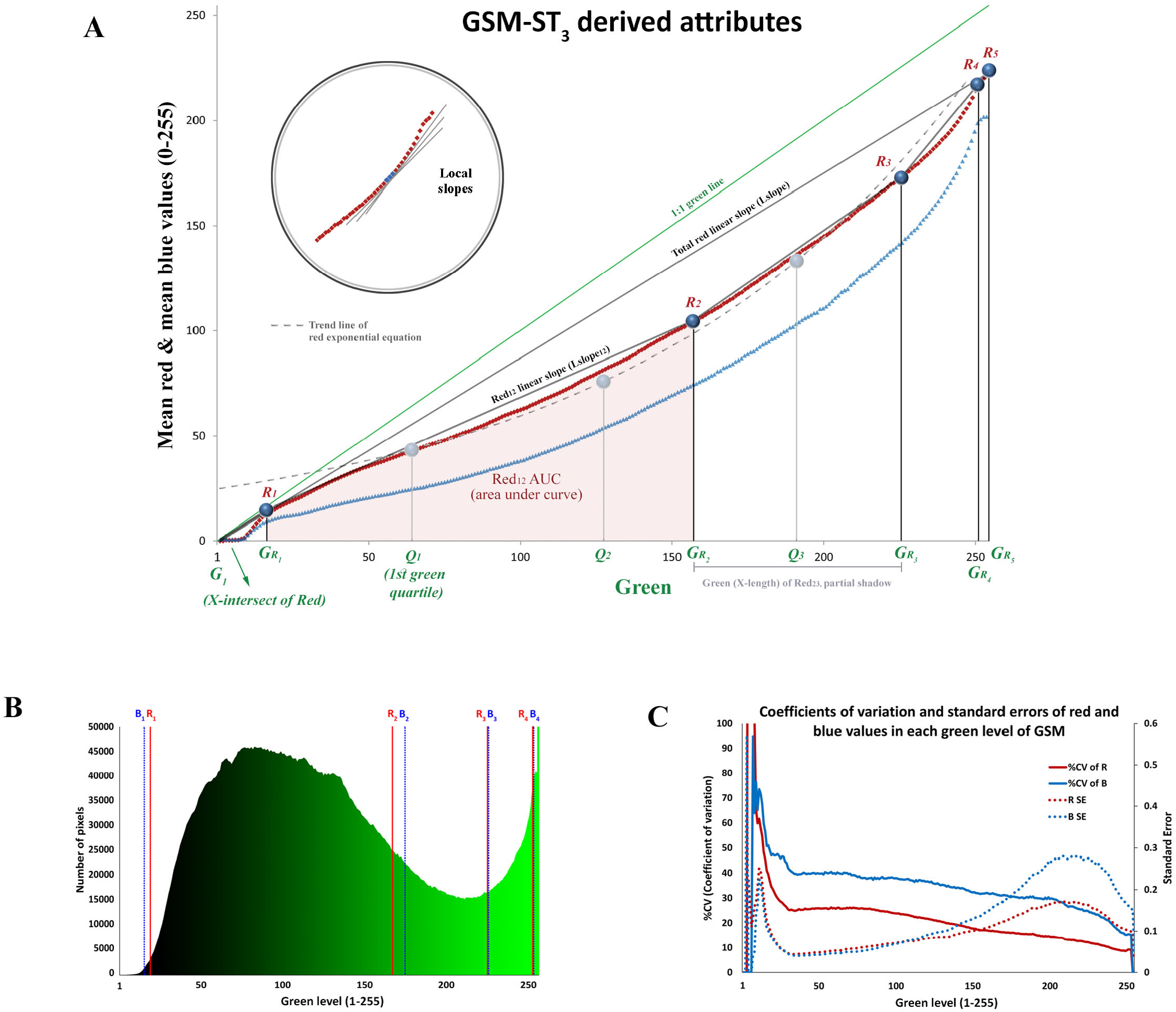
GSM-curve characteristics. **(A)** Schematic illustration of GSM graph and some attributes defined based on segmentation type 3 (ST_3_); which may be utilized for datamining evaluations parallel to the two approaches described in the present study. Examples of these ST_3_-attributes include under- and between curve areas, linear and local slopes ofthe curves, and level ofred or blue color at the certain quartiles (or percentiles) ofthe horizontal axis length. **(B)** Distribution ofimage pixels participated in drawing GSM graph. R_1_ and/or B_1_ to R_4_ and/or B_4_ are the thresholds ofsegmentation type 3. **(C)** Indices ofdispersion calculated for GSM graph. As described in the manuscript, the GSM curves are formed based on the mean values ofred or blue colors ofthe vegetation part. Here, the coefficients of variation (%) and standard errors for red and blue colors in each green level are shown.

**Supplementary Figure 4.**
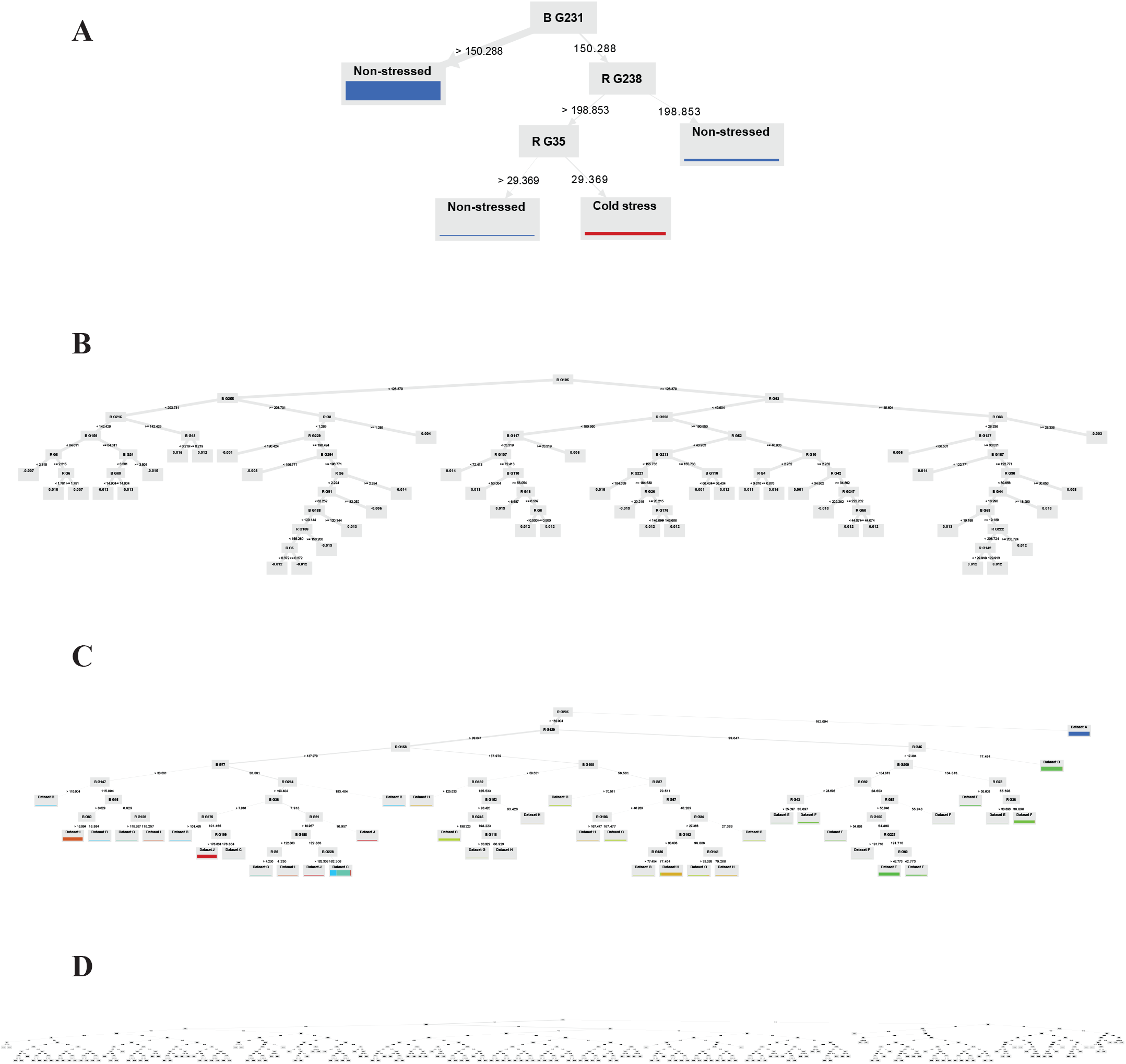
Example outputs of datamining models used for prediction of various traits based on GSM-ST2 attributes. The example trees are selected from the sets of Random Forest (RF) and Gradient Boosted Tree (GBT) models used for: **(A)** distinguishing between the cold-stressed and non-stressed canopies (RF); **(B)** classification of canopies based on whether they were imaged before or after irrigation (irrigation event analysis; GBT); **(C)** classification of canopies based on the dataset they were belonged to (i.e. imaging date; RF); and **(D)** prediction of canopy temperature (°C; RF). In all of the parts, the most complete combination of the relevant datasets (i.e. dataset “Total”) is used both for training and testing the models. R G_i_ and B G_i_ are GSM-ST_2_ attributes, which show the points on the red and blue curves (mean red and mean blue values) at the green level of *i*, respectively.

**Supplementary Figure 5.**
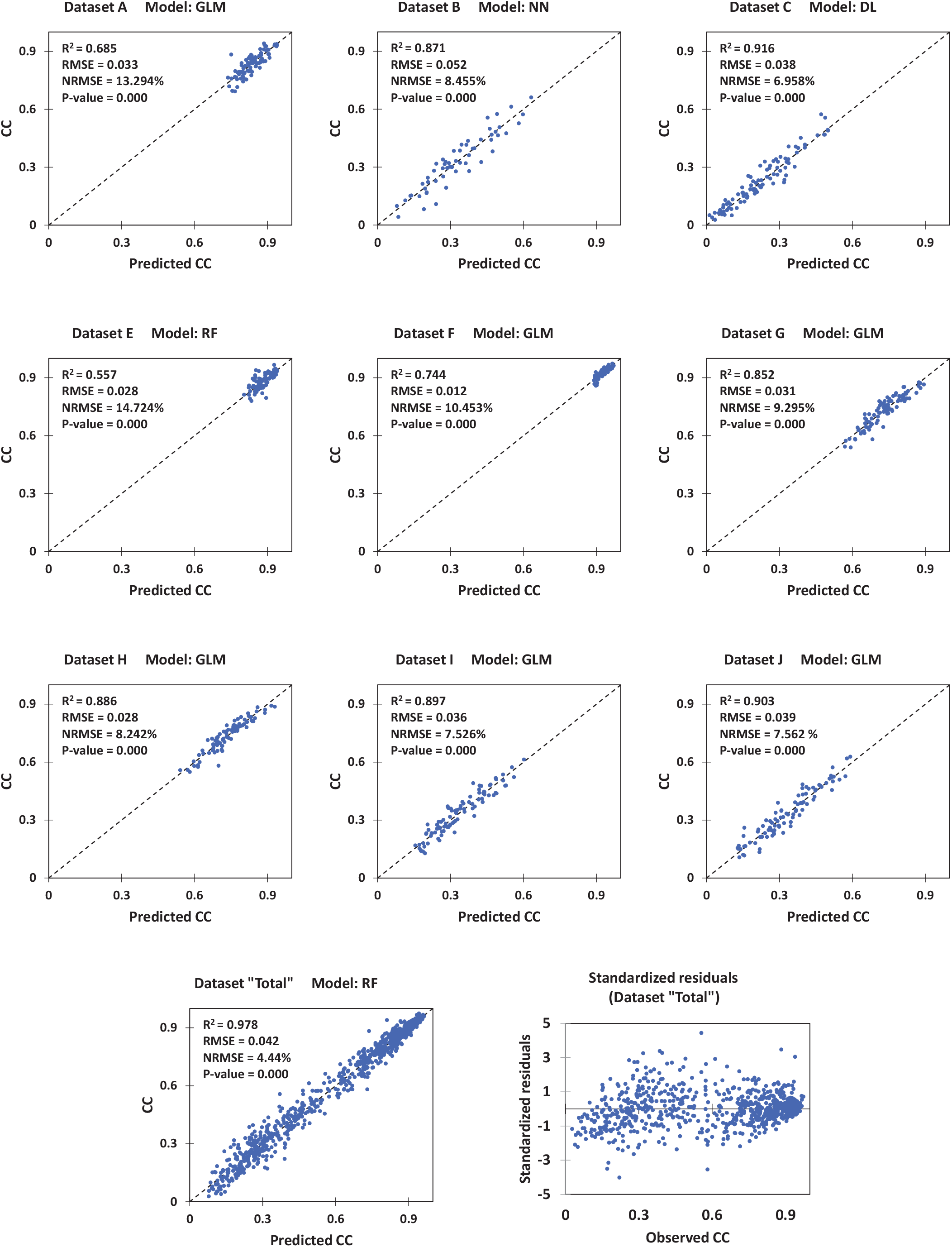
Cross-validation for prediction of canopy coverage (CC) by datamining models using coefficients of GSM exponential equations. The dataset “Total” is combination of all other datasets. For each dataset, results of the model with the best performance are shown. DL: Deep Learning; GLM: General Linear Model; NN: Neural Network; RF: Random Forest. NRMSE: normalized root mean square error (calculated by dividing the RMSE by the range of observed values).

**Supplementary Figure 6.**
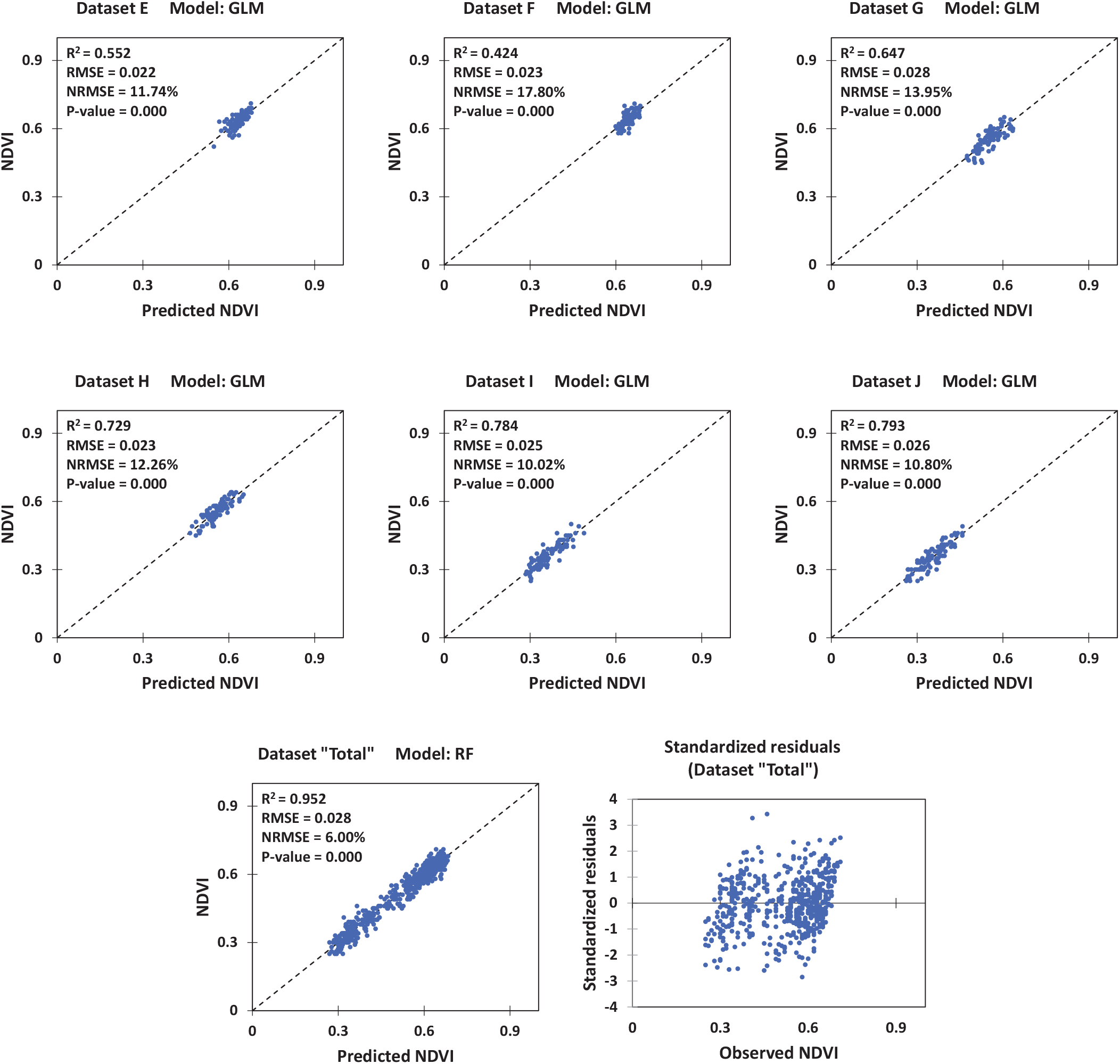
Cross-validation for prediction of Normalized Difference Vegetation Index (NDVI) by datamining models using coefficients of GSM exponential equations. The dataset “Total” is combination of all other datasets. For each dataset, results of the model with the best performance are shown. GLM: General Linear Model; RF: Random Forest. NRMSE: normalized root mean square error (calculated by dividing the RMSE by the range of observed values).

**Supplementary Figure 7.**
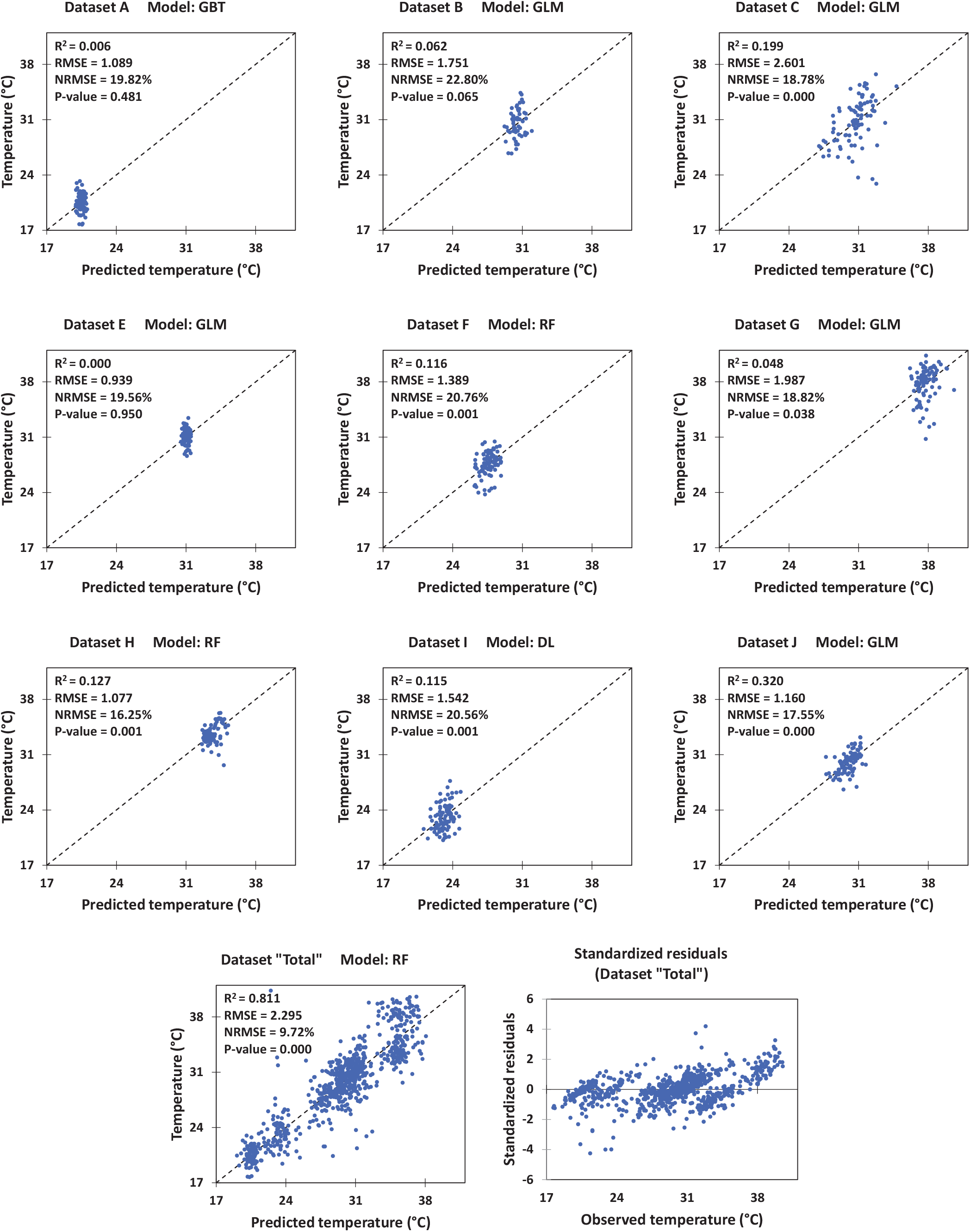
Cross-validation for prediction of canopy temperature (°C) by datamining models using coefficients of GSM exponential equations. The dataset “Total” is combination of all other datasets. For each dataset, results of the model with the best performance are shown. DL: Deep Learning; GBT: Gradient Boosted Trees; GLM: General Linear Model; RF: Random Forest. NRMSE: normalized root mean square error (calculated by dividing the RMSE by the range of observed values).

**Supplementary Figure 8.**
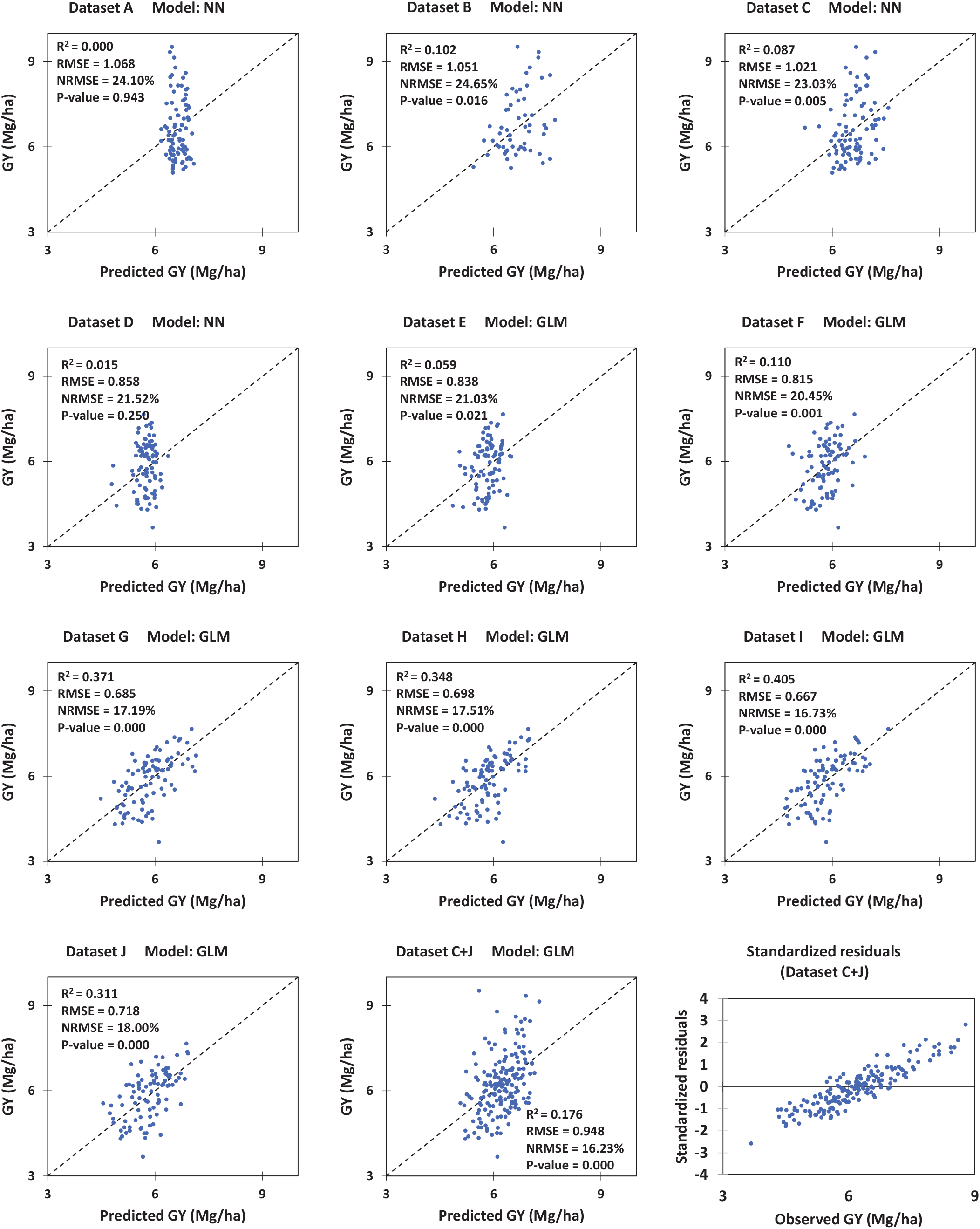
Cross-validation for prediction of grain yield (GY) by datamining models using coefficients of GSM exponential equations. As datasets C and J (included the images taken at the last imaging dates in the 1^st^ and 2^nd^ seasons) had relatively the highest relationships with GY, their combination was also tested as a single dataset named “C+J”. For each dataset, results of the model with the best performance are shown. GLM: General Linear Model; NN: Neural Network. NRMSE: normalized root mean square error (calculated by dividing the RMSE by the range of observed values).

**Supplementary Figure 9.**
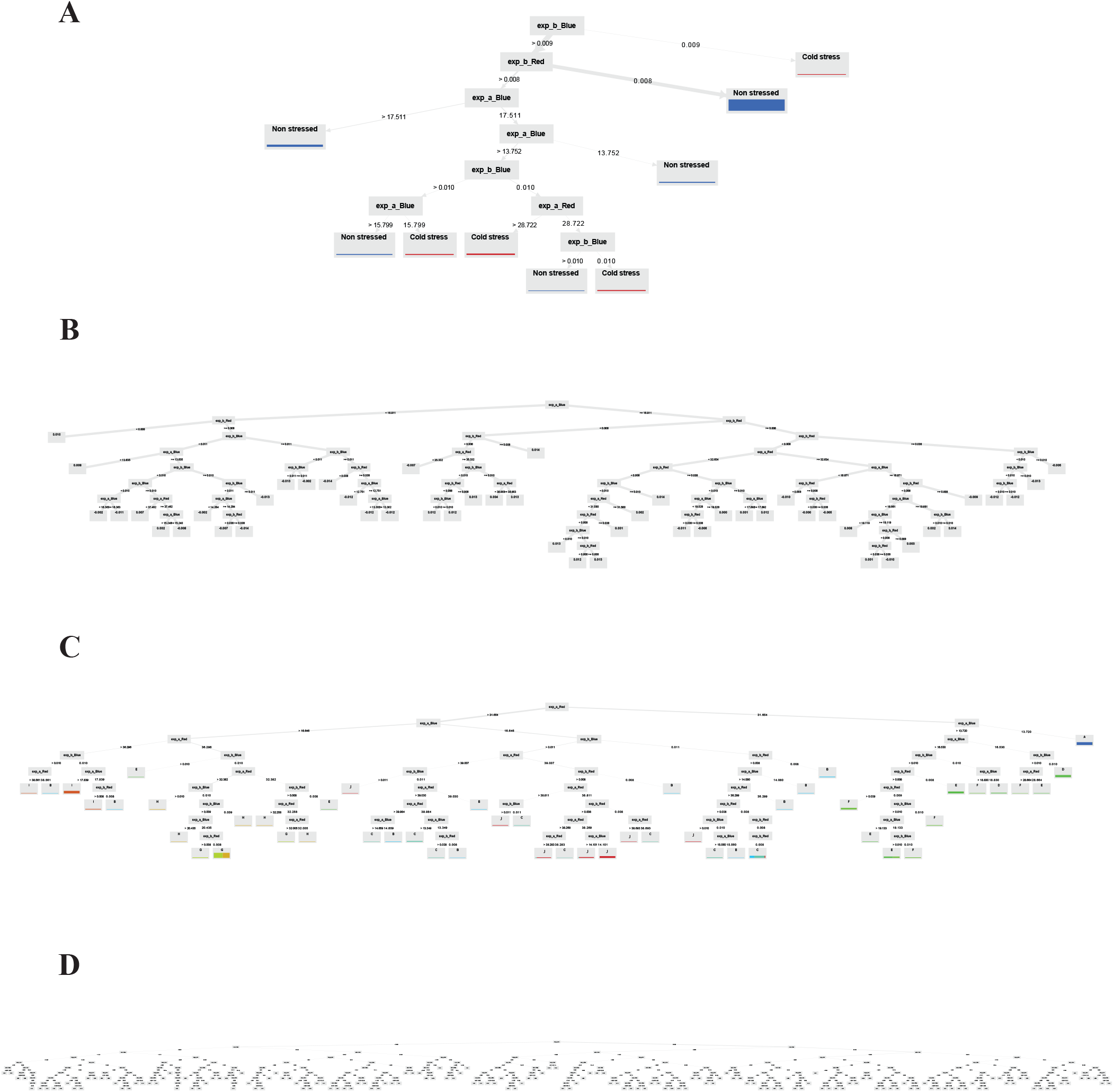
Examples outputs of datamining models used for prediction of various traits based on the coefficients of GSM exponential equations. The example trees are selected from the sets of Random Forest (RF) and Gradient Boosted Tree (GBT) models used for: **(A)** distinguishing between the cold-stressed and non-stressed canopies (RF); **(B)** classification of canopies based on whether they were imaged before or after irrigation (irrigation event analysis; GBT); **(C)** classification of canopies based on the dataset they were belonged to (i.e. imaging date; RF); and **(D)** prediction of NDVI (Normilized Difference Vegetation Index; RF). In all of the parts, the most complete combination of the relevant datasets (i.e. dataset “Total”) is used both for training and testing the models. *Red*: GSM red trend, *Blue*: GSM blue trend, exp-*a* and -*b*: coefficients *a* and *b* of the GSM exponential equations.

**Supplementary Figure 10.**
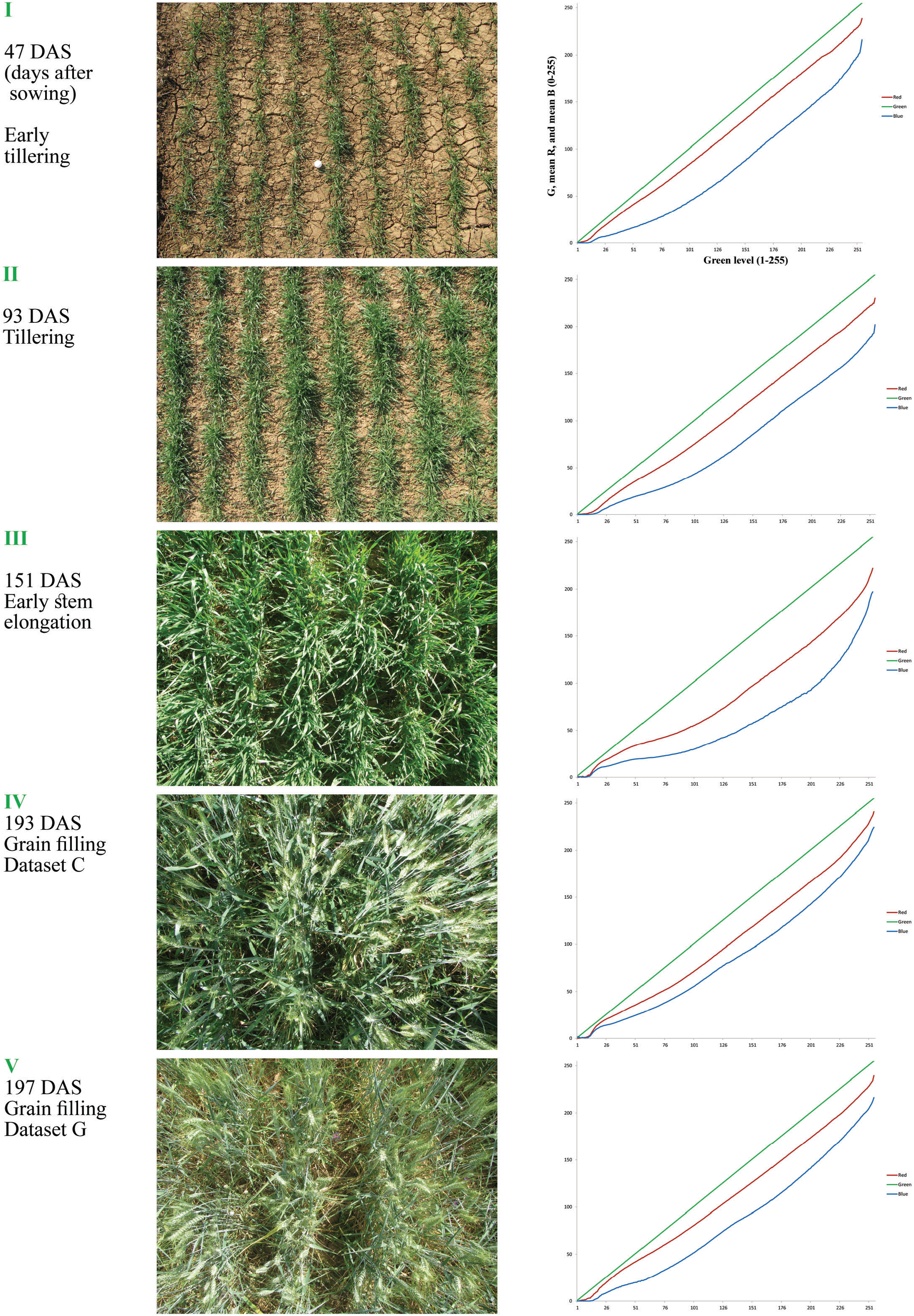
Effect of growth stage (phenology) on GSM graph in a green monoculture canopy. Consider the variations of the red and blue curves relative to green line, which in the most dense canopy at the stage (III) show the highest degrees of curvature.

**Supplementary Figure 11.**
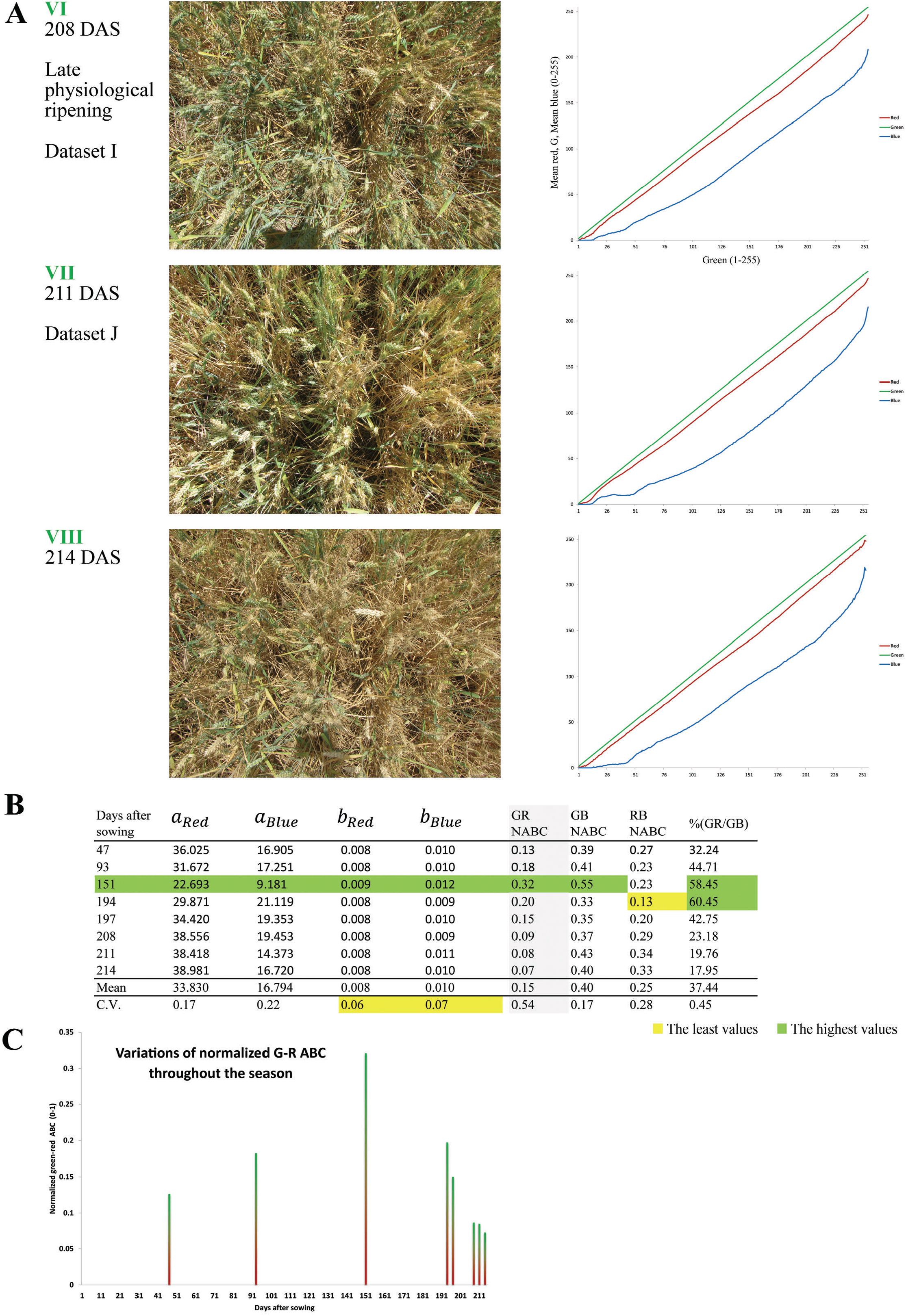
GSM graph ofa canopy at different degrees ofripening. **(A)** GSM seems to be capable to track the canopy status even in a semi-dried stand. Consider the shape ofthe red curve which is approached to the green line, late in the season. It is notable that only green pixels are included in the GSM analyses. **(B)** Various phenological stages are compared using the exponential equations and ST_3_-derived curve attributes. R, G, and Bare levels of red, green, and blue colors; NABC is normalized area between curves, which is calculated by dividing each ABC value to the total area under green line. **(C)** GR-NABC is recommendable for determining the canopy growth stage, and tracking ofripening.

**Supplementary Figure 12.**
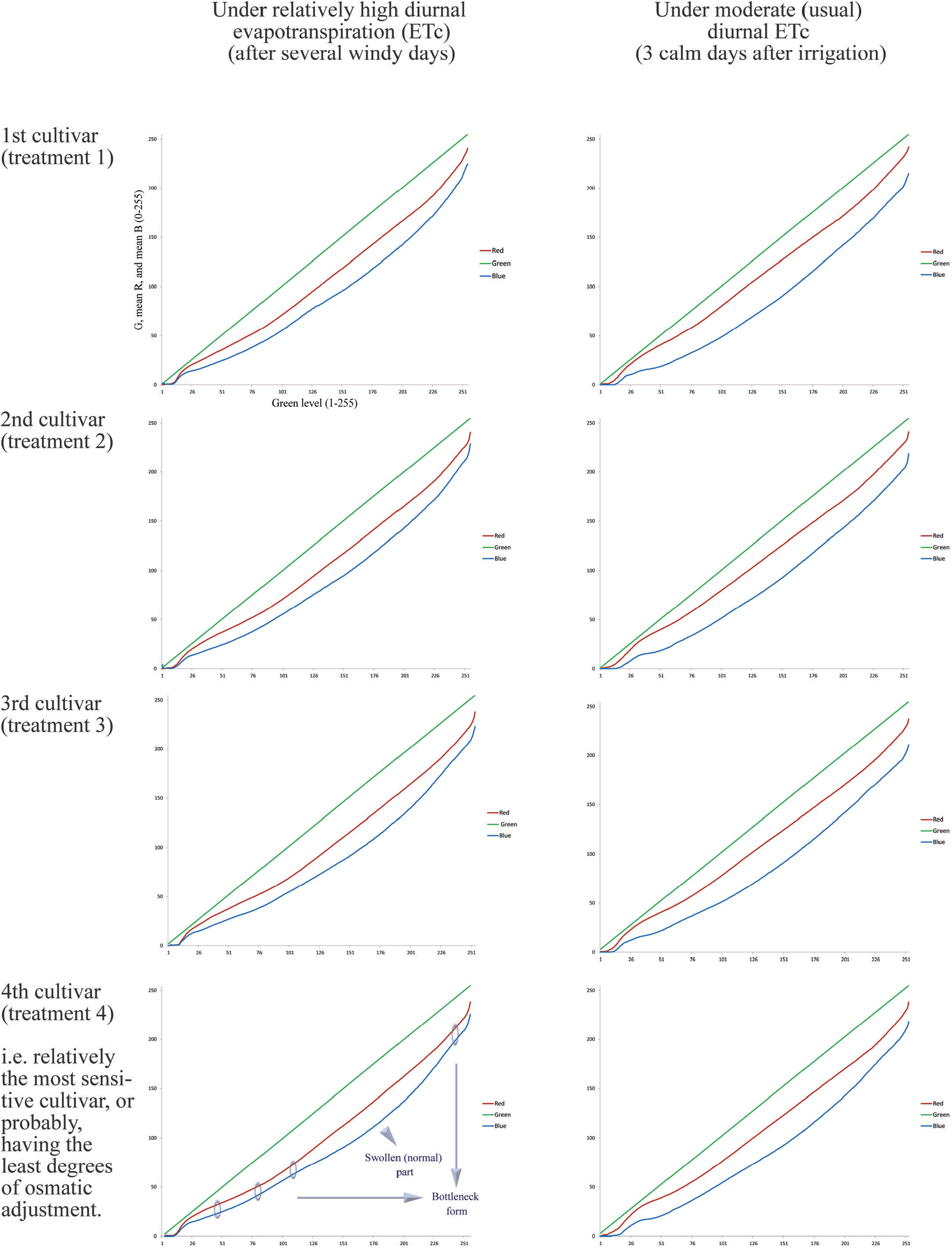
Effect of water stress on the shape of GSM graph in the canopies of 4 wheat cultivars. It seems that the overall result ofvariations in red and blue curves, lead to bottleneck and swollen forms, e.g. due to higher reflection ofblue light under complete shadow. It is noteworthy, that the reflection pattern is expected to be influenced by the interaction between the intensity of stressful condition and degree of osmotic adjustment, which also may result to different responses among cultivars.

**Supplementary Figure 13.**
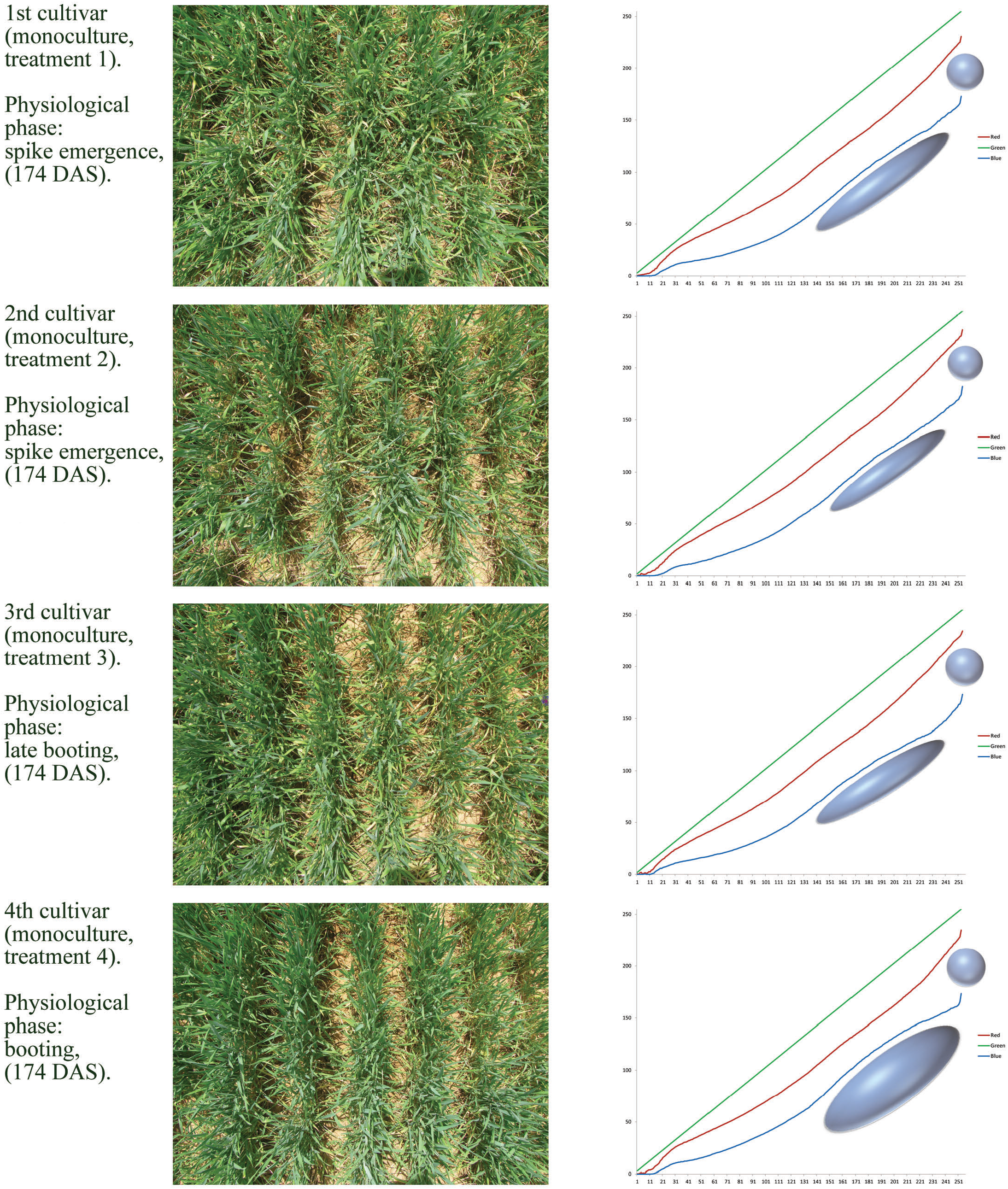
Images and GSM graphs ofthe canopies of4 wheat cultivars, 4 days after a moderate cold stress. The most obvious impact was on the blue curve, which has resulted to a diversion from the normal trend, i.e. an increase in the level ofmean blue values at the middle of the curve (green points either exposed to direct sunlight, or positioned under partial shadow, see the manuscript). Moreover, compared to the curves of non-stress conditions, there is a more considerable difference between the maximum values of red and blue (near 255, in full sunlight). The interferences of cold stress have shown by ellipsoids and spheres of different sizes (which may be related to different tolerances of the cultivars).

## Revision summary

This version of the manuscript has been revised to update the following:

i. GY and GN predictions: it is a pity for us to report an unwanted mistake which has occurred in datamining analyses of the previous version. Very recently, while we were working on extension of GSM for other physiological approaches, it came in to our notice that the calculations related to prediction of GY and GN using GSM outputs were not correct as such! Indeed, due to a mistake in designing the computation processes in the environment of RapidMiner software, a single dataset was used for both training (construction) and validation of the data mining models. As the result, the accuracies reported for the prediction of GY and GN in the 2^nd^ and 3^rd^ computation approaches were highly overestimated. Here, an operator (i.e. Split Data) was used to split the dataset randomly into two 45 subsets for training and validation of datamining models. Unfortunately, one of the connections from the output terminals of this operator was missed. After finding the mistake, we re-ran the data mining models with two appropriately distinct datasets (each included data of 45 plots) and found that the accuracies were much lower than what has been reported in the manuscript (even lower than 50%). This suggests that at least in the conditions of the present experiment (or having only 90 observations), GY and GN could not be predicted accurately using the reported method. So, GN analysis is excluded from the new version. We should emphasize that the calculations and procedures reported in other parts of the manuscript are all correct and were not affected by this unwanted mistake.
ii. A practical shift in datamining approaches: although the previous datamining approach (which was developed based on ST_3_) was beneficial and efficient for other purposes, its adoption in datamining analyses may require recalculation of variables used in the algorithm in response to dynamics of canopy or environmental conditions. Therefore, in the new version, the ST_3_-based approach was replaced with two other parallel approaches:

a. Based on the ST_2_-derived attributes (up to 510 attributes), and
b. Based on the coefficients of exponential equations fitted to the GSM red and blue curves (4 attributes). As shown by results, both of the recent approaches are recommendable and may be utilized almost in every conditions without additional calculation or the need for resetting variables.
iii. Evaluation of new quantitative and qualitative characteristics of canopy: in the new version, potential of GSM in prediction of new traits is evaluated.
iv. Processing a bigger dataset: despite the previous version in which 90 images were analyzed, in the current version a total number of 866 images taken in 10 imaging dates were used for datamining analyses.
v. Sharing the MATLAB code: now, a user-friendly version of the exclusive MATLAB code used in the present study is published as a reproducible compute capsule (i.e. Canopy GSM) on Code Ocean, which is available at: https://doi.org/10.24433/CO.4355649.v1

